# EMT-induced cell mechanical changes enhance mitotic rounding strength

**DOI:** 10.1101/598052

**Authors:** Kamran Hosseini, Anna Taubenberger, Carsten Werner, Elisabeth Fischer-Friedrich

## Abstract

To undergo mitosis successfully, most animal cells need to acquire a round shape to provide space for the mitotic spindle. This mitotic rounding relies on mechanical deformation of surrounding tissue and is driven by forces emanating from actomyosin contractility. Cancer cells are able to maintain successful mitosis in mechanically challenging environments such as the increasingly crowded environment of a growing tumor, thus, suggesting an enhanced ability of mitotic rounding in cancer. Here, we show that epithelial mesenchymal transition (EMT), a hallmark of cancer progression and metastasis, gives rise to cell-mechanical changes in breast epithelial cells. These changes are opposite in interphase and mitosis and correspond to an enhanced mitotic rounding strength. Furthermore, we show that cell-mechanical changes correlate with a strong EMT-induced change in the activity of Rho GTPases RhoA and Rac1. Accordingly, we find that Rac1 inhibition rescues the EMT-induced cortex-mechanical phenotype. Our findings hint at a new role of EMT in successful mitotic rounding and division in mechanically confined environments such as a growing tumor.

## Introduction

Most animal cells adopt an approximately spherical shape when entering mitosis^1^. This process has been termed mitotic rounding. It ensures the correct morphogenesis of the mitotic spindle and, in turn, successful cell division^2–5^. When cells acquire a round shape at the entry of mitosis, they need to mechanically deform the surrounding tissue to do so (Figure 1). Previous studies suggest that the forces necessary for this deformation emerge from the contractility of the mitotic actin cortex^1,3,5–11^. In fact, at the onset of mitosis, cortical contractility was found to be upregulated giving rise to an increased cell surface tension which drives the mitotic cell into a spherical shape^8,9^. This physical picture is consistent with reports that mitotic rounding relies on RhoA^7,12,13^ – a major actomyosin regulator in the cell.

**Figure 1:**
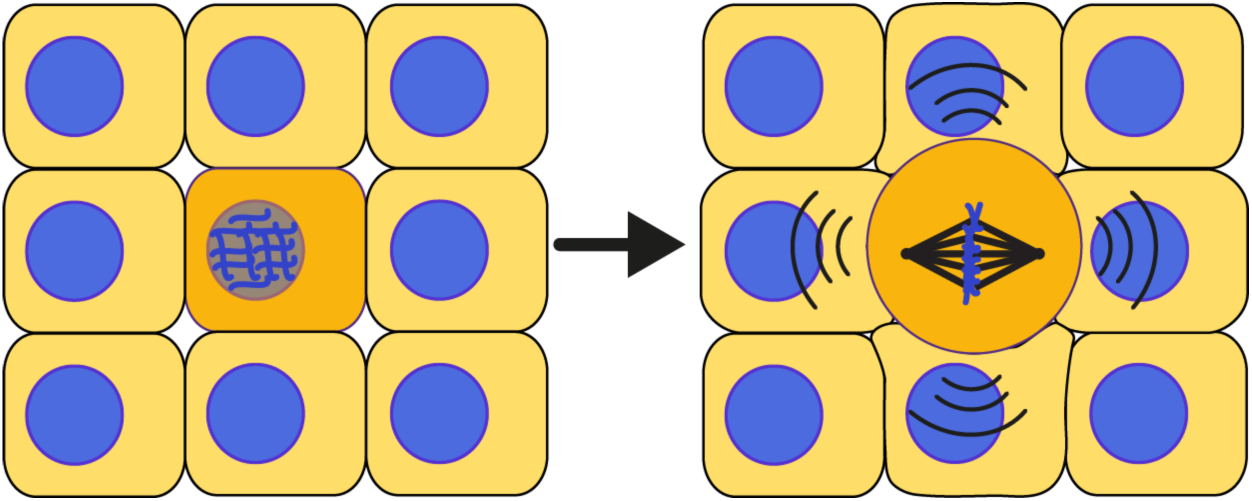
Schematic of mitotic rounding in a tissue: At the onset of mitosis, cells need to deform their surrounding when acquiring a spherical shape in mitosis. Mechanical forces for rounding emerge from actomyosin contractility of the mitotic cell cortex.

In a growing tumor, an increasing cell density generates a compressive mechanical stress which would likely lead to an increasing mechanical obstacle for mitotic rounding. Indeed, mechanical confinement or external pressure have been shown to hamper cell proliferation in tumor spheroids^14–18^. Thus, it has been hypothesized that the actin cortex of cancer cells exhibits oncogenic adaptations that allow for ongoing mitotic rounding and division inside tumors^19^. In fact, it was shown that the human oncogene Ect2 contributes to mitotic rounding through RhoA activation^7,10^ and that Ras overexpression promotes mitotic rounding^20^.

Epithelial-mesenchymal transition (EMT) is a cellular transformation in which epithelial cells loose epithelial polarity and intercellular adhesiveness gaining migratory potential^21–23^. EMT, a hallmark in cancer progression, is commonly linked to early steps in metastasis promoting cancer cell invasiveness. Moreover, EMT was connected to cancer stem cells and the outgrowth of secondary tumors^21–23^, suggesting that EMT may also be important for cell proliferation in a tumor.

In this work, we test the hypothesis that EMT enhances mitotic rounding strength. To assess mitotic rounding strength, we measure the mechanical properties of the actin cortex in mitosis, in particular cortex stiffness and contractility before and after EMT. Furthermore, we also determine mechanical changes of the actin cortex of interphase cells upon EMT; mechanics of interphase cells may critically influence mitotic rounding as interphase cells are a major constituent of the surrounding of a mitotic cell which needs to be deformed in the process of rounding (Figure 1). For our cortex-mechanical measurements, we use an established dynamic cell confinement assay based on atomic force microscopy^9,24^. We report cortex-mechanical changes upon EMT that are opposite in interphase and mitosis. They are accompanied by a strong change in the activity of the actomyosin master regulators Rac1 and RhoA. Complementary, we characterize cortex-mechanical changes induced by Rac1 and RhoA signaling. In particular, we show that Rac1 inhibition restores epithelial cortex mechanics in post-EMT cells. Furthermore, we give evidence that EMT, as well as Rac1 activity changes induce actual changes in mitotic rounding in spheroids embedded in mechanically confining, covalently crosslinked hydrogels^25^.

## Results

### Breast epithelial cells undergo EMT upon drug treatment

For our study, we use two established human breast epithelial cell lines MCF-10A and MCF-7. MCF-10A are pre-neoplastic cells which serve as a model for mammary epithelial cells in healthy conditions, MCF-7 are cancerous cells derived from a metastatic mammary carcinoma. We induce EMT in MCF-7 and MCF-10A breast epithelial cells respectively through established methods by incubation with phorbol ester (TPA)^26^ or TGF-*β*1^27–29^, respectively (Figure S1, Materials and Methods). Accordingly, both cell lines showed a loss of the epithelial marker E-cadherin and an increase in mesenchymal markers Vimentin and N-Cadherin after treatment according to Western blot quantification (Figure S1).

TPA-treatment enhances proliferation and migration in MCF-7 cells corresponding to a more aggressive cancer cell phenotype^26^. In addition, TGF-*β*1-induced transformation of MCF-10A was shown to enhance cell migration^30^ but to inhibit cell proliferation in 2D culture (Figure S1). This proliferation effect is reversed by oncogenic signaling^31^.

In the following, cells that underwent EMT will be denoted as modMCF-7 and modMCF-10A, respectively.

### Measurement of actin cortex mechanics

We probed actin cortex mechanics in mitotic and interphase cells through a previously established cell confinement setup based on atomic force microscopy (AFM). In this assay, initially round cells are dynamically confined between two parallel plates using a wedged cantilever^32–35^ (Figure 2a,b, Materials and Methods). This method was shown to measure the contractility and the viscoelastic stiffness of the actin cortex^33,34^. Note that our assay relies on an initially spherical cell shape. Therefore, interphase cells are sampled in a state of suspension which also enhances comparability to measurements of rounded mitotic cells.

**Figure 2:**
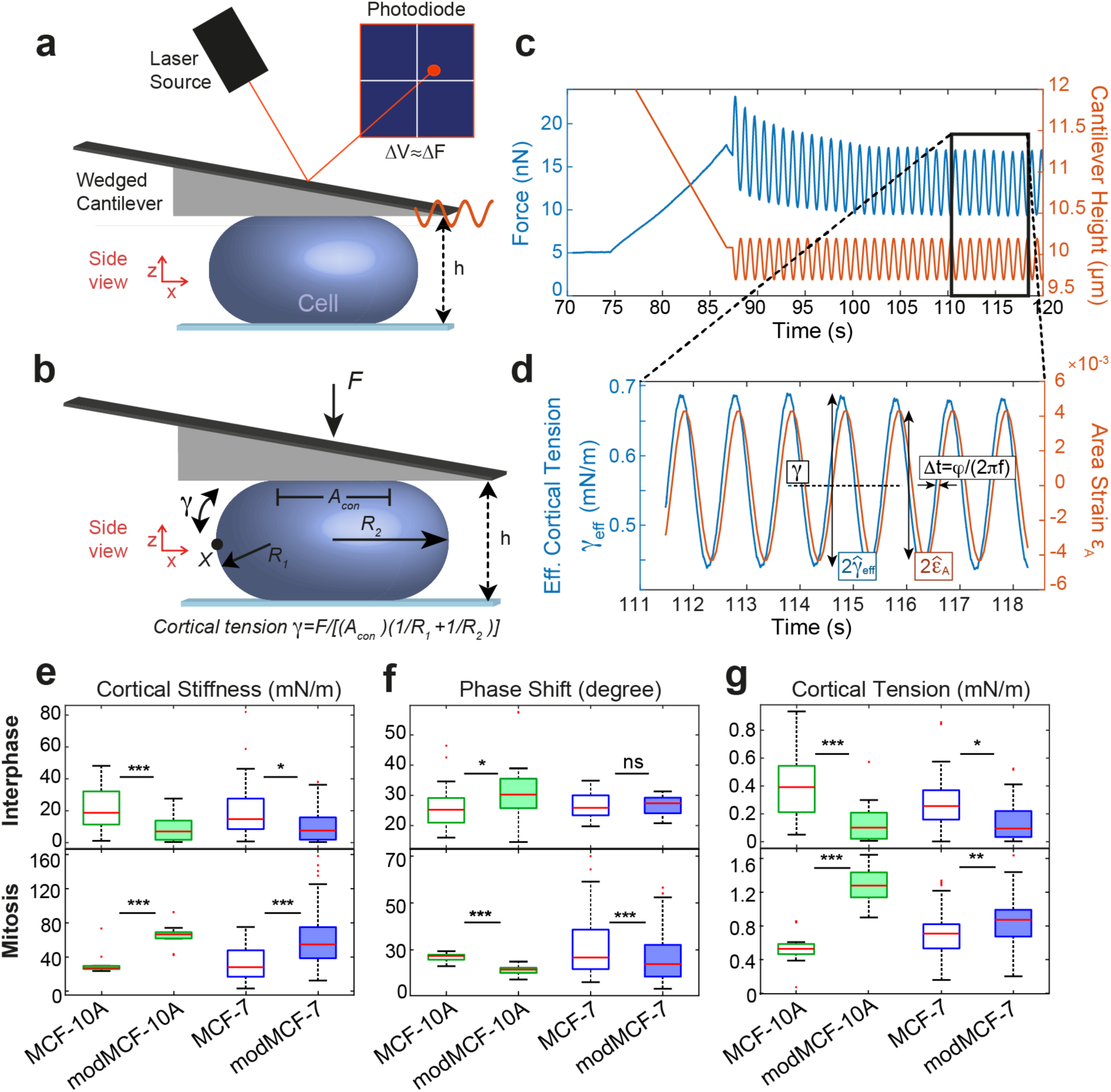
Mechanical phenotyping of the actin cortex of MCF-7 and MCF-10A before and after EMT measured by dynamic AFM cell confinement. **a**, Schematic of measurement setup: initially round cells (mitotic cells or suspended interphase cells) are confined by the tip of a wedged AFM cantilever and oscillatory height oscillations are imposed. **b**, Mechanical and geometrical parameters of a measured cell used for data analysis. **c**, Exemplary AFM readout during cell confinement and subsequent oscillatory cell height modulation: force (blue), cantilever height (orange). **d**, Effective time-dependent cortical tension estimate (blue) and relative cell surface area change (orange) from data analysis of the indicated time window in c. Mechanical parameters are determined as follows: **cortex stiffness** K is calculated as 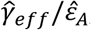, **phase shift** is obtained as φ = (2πf)Δt, **cortical tension** is calculated as time average of γ_eff_ (dashed black line). The corresponding complex elastic modulus of the cortex is K^*^ = Ke^iφ^. **e-g**, Cortical stiffness (**e**), phase shift (**f**) and cortical tension (**g**) measured for suspended interphase cells (top row) and cells in mitotic arrest (bottom row) before and after EMT. (Post-EMT cells are referred to as modMCF-7 and mod-MCF-10A, respectively. Number of cells measured: Interphase: MCF-7 n=27, modMCF-7 n=28, MCF-10A n=27, modMCF-10A n=31, Mitosis: MCF-7 n=34, modMCF-7 n=45, MCF-10A n=12, modMCF-10A n=12. Measurements are representative for three independent experiments. ^ns^p> 0.05, *p < 0.05, **p < 0.01, ***p < 0.001).

During measurements, we confined cells first to the desired confinement height and then imposed small cell height oscillations at a frequency of f=1 Hz, which triggered a cellular force response measured by the AFM (Figure 2c). Using previous insight^34^, we calculated a normalized change of cortical surface area (area strain) and an effective cortical tension from the output height and force signal (Figure 2d, Materials and Methods). These were, in turn, used to extract a complex elastic modulus of cortex mechanics at the chosen oscillation frequency (Materials and Methods). In the following, we will refer to the absolute value of the complex elastic modulus as cortical stiffness. The complex phase of this modulus will be referred to as phase shift. This phase shift indicates the nature of the cortex material: a phase shift of 0 degree corresponds to a solid material, a phase shift of 90 degrees corresponds to a liquid material, while values in between indicate an intermediate viscoelastic behavior^34^. We thereby obtain three mechanical parameters of the cortex from the measurement of each cell: cortical stiffness, phase shift and cortical tension. These three parameters provide a detailed mechanical fingerprint of the cortical architecture of the cell.

### EMT changes cortex mechanics and composition

We tested how the mechanics of the actin cortex changes through EMT to assess the associated change in mitotic rounding strength. For this purpose, we applied our cell-mechanical assay to cells before and after EMT induction (Figure 2e-g). Measurements on interphase cells show a striking decrease in cortical tension and stiffness following EMT induction (Figure 2e-g, upper row). In addition, we find that the measured phase shift increased in MCF-10A but did not change significantly in MCF-7 (Figure 2f, upper row).

In a second step, we examined the mechanical properties of the cell cortex in mitotic cells before and after EMT. For mechanical measurements, cells were pharmacologically arrested in mitosis by addition of S-trityl-L-cysteine (STC) to enrich mitotic cells and prevent cell-cycle related mechanical changes (see Materials and Methods)^7,35,36^. STC-treated cells were previously shown to exhibit cell-mechanical properties comparable to cells in metaphase of mitosis^8,34^. To our surprise, we now observed a change opposite to our finding in interphase cells: cortical tensions and cortical stiffness were clearly increased in both cell lines while phase shifts went down (Figure 2e-g, lower row). In summary, our measurements show that the post-EMT mitotic cortex is stiffer, more contractile and more solid-like for both cell lines.

A more contractile mitotic cortex in combination with a softened actin cortex in interphase suggests an enhanced ability to undergo mitotic rounding post-EMT. Observed mechanical changes were substantial for both cell lines, but in general stronger for MCF-10A (Figure 2). We were interested in the molecular origin of the observed EMT-induced cortex-mechanical changes. Therefore, we investigated changes of actin and myosin levels in MCF-7 cells upon EMT. First, we quantified total levels of actin and myosin by immunoblotting before and after EMT in asynchronous cell populations (mainly interphase) and in mitotic cells (Figure 3a). While levels of actin are unaffected by EMT, we find that myosin levels show an EMT-induced rise in mitosis (Figure 3a). In a complementary experiment, we quantified the cortex-to-cytoplasm ratio of actin and myosin before and after EMT (Figure 3b). To this end, we transfected MCF-7 cells in pre- and post-EMT conditions with plasmids expressing either actin (ACTB) fused to mCherry or myosin light chain 9 (Myl9) fused to mApple (see Materials and Methods). Subsequently, we imaged the equatorial cross-section of these cells and quantified the ratio of cortical versus cytoplasmic fluorescence intensity (Figure 3b, S3a-d, Materials and Methods). We found that the respective cortex-to-cytoplasm ratio of myosin decreases in interphase cells upon EMT (Figure 3b). By contrast, in mitosis, we did not observe significant changes of myosin distributions (Figure 3b).

**Figure 3:**
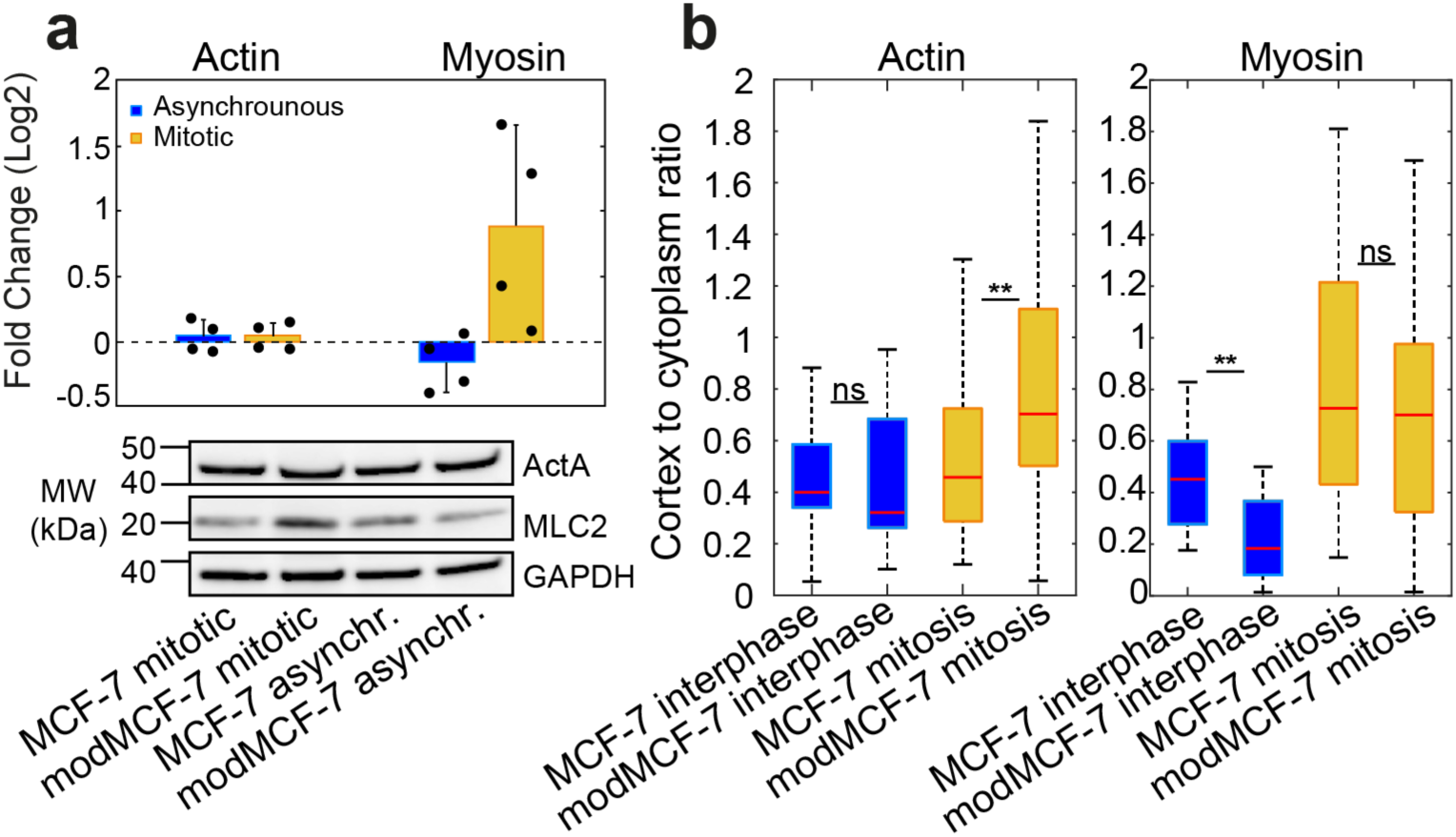
Actin and myosin changes in the cortex upon EMT. **a**, Relative changes of myosin and actin expression from Western blots. Top panel: Quantification of relative changes, individual data points are shown with black dots. Bottom panel: Western blots showing actin (top row) and myosin (second row) abundancy before and after EMT in asynchronous cell populations (mainly interphase) and mitotic cells. **b**, Ratio of cortical versus cytoplasmic actin (left panel) or myosin (right panel) in suspended interphase cells and STC-arrested mitotic cells, pre- and post-EMT. (Post-EMT cells are referred to as modMCF-7. Number of cells measured: Panel **a**: n=4. Panel **b**: Actin: MCF-7 interphase n=18, modMCF-7 interphase n=16, MCF-7 mitotic n=35, modMCF-7 mitotic n=33. Myosin: MCF-7 interphase n=17, modMCF-7 interphase n=15, MCF-7 mitotic n=17, modMCF-7 mitotic n=17. Measurements are representative for at least two independent experiments. ^ns^p> 0.05, *p < 0.05, **p < 0.01, ***p < 0.001).

Considering both experimental findings together (immunoblotting and cortex-to-cytoplasm ratios), our data suggest that the amount of cortex-associated myosin increases in mitosis but decreases in interphase cells upon EMT (Figure 3a,b). Therefore, levels of cortex-associated myosin exhibit positive correlation with reported changes of cortical tension and stiffness in interphase and mitosis. Changes of cortex-associated myosin may thus provide a simple mechanistic explanation for EMT-induced cortex-mechanical trends.

Furthermore, we found no changes of the cortex-to-cytoplasm ratio of actin upon EMT in interphase (Figure 3b). However, we observed a significant increase of this ratio in mitosis upon EMT (Figure 3b, Figure S3d). This puts forward the option of an EMT-induced increase in actin cortex density and/or actin cortex thickness in mitosis. To rule out one or the other scenario, we estimated cortical thickness in mitotic cells (Figure S3e,f). Our findings suggest either no thickness change or slightly diminished cortical thickness post-EMT, thus, favouring the scenario of an actin cortex density increase.

In addition, we monitored actin turnover at the cortex of MCF-7 cells before and after EMT (Figure S3g). Particularly in post-EMT mitosis, we find a trend towards faster actin turnover further hinting at modified dynamics and structure of cortical actin in post-EMT cells.

### EMT upregulates activity of Rac1 and downregulates activity of RhoA

Since Rho-GTPases are known to be key regulators of the actin cortex, we tested the changes in activity of the Rho-GTPases RhoA, Rac1 and Cdc42 by performing pull-down assays of active GTP-bound Rho-GTPases from cell lysates using beads (see Materials and Methods). For Cdc42, we did not find obvious changes upon EMT in an asynchronous cell population (mainly interphase cells) or in isolated mitotic cells (Figure S4). However, we observed a drastic increase of active Rac1 accompanied by a strong decrease of active RhoA upon EMT in both mitotic cells and for asynchronous cell populations (Figure 4a,b). EMT-induced activity reduction of RhoA was more striking for asynchronous cell populations than for mitotic cells, indicating that RhoA was still activated in mitosis post-EMT.

**Figure 4:**
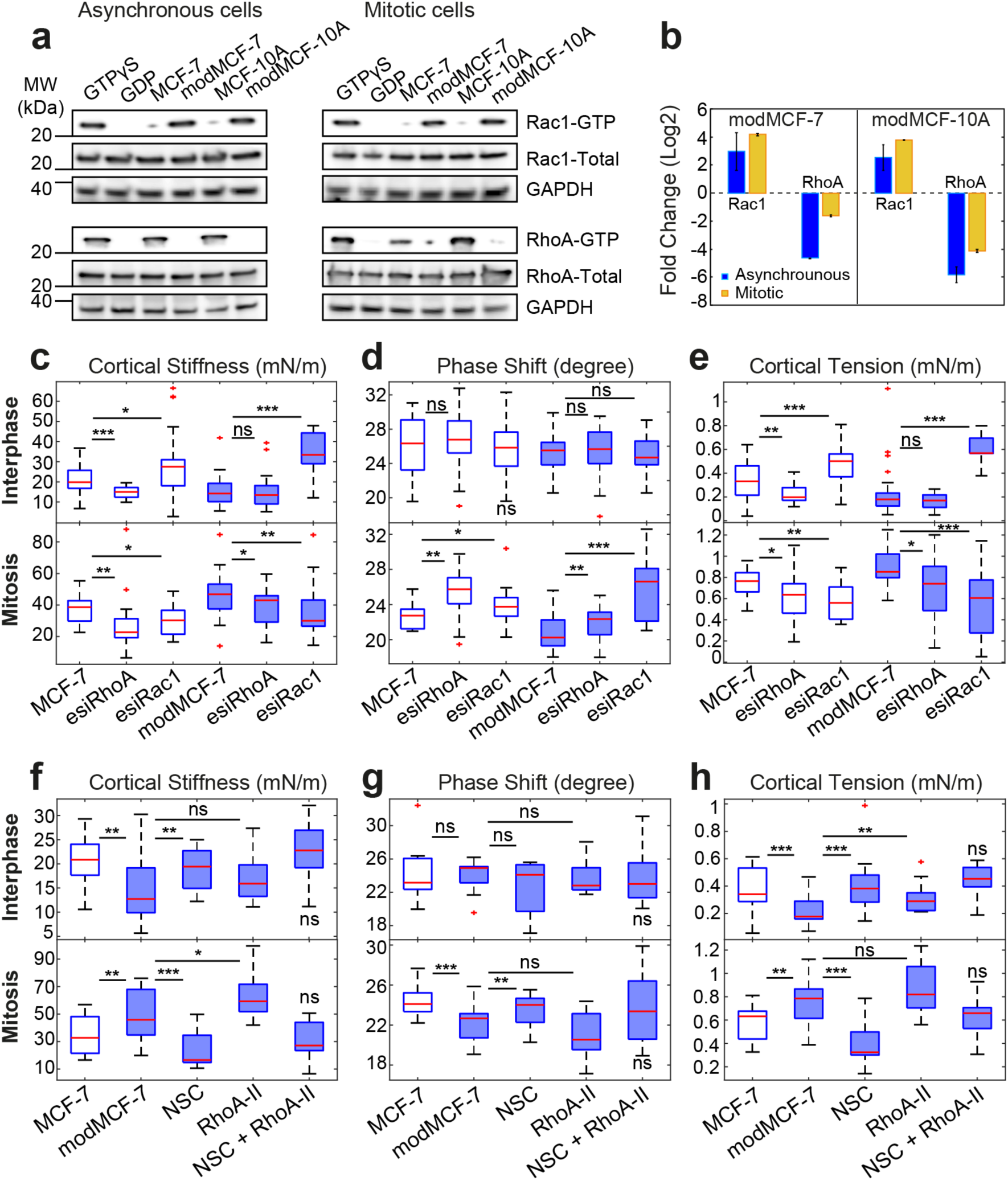
Activity changes of Rac1 and RhoA upon EMT and related changes in cortical mechanics. a,b) Rac1 and RhoA activity before and after EMT. **a**, Western blots showing GTP-bound Rac1 (top row) and RhoA (third row) before and after EMT. **b**, Quantification of relative changes of Rac1-GTP and RhoA-GTP from Western blots in **a**. Quantifications were normalised by GAPDH band. **c-e**, Effect of Rac1 and RhoA knock-down on cortex mechanics in MCF-7 and modMCF-7 (post-EMT) in suspended interphase MCF-7 cells (top row) and MCF-7 cells in mitotic arrest (bottom row). **f-h**, Effect of RhoA activation upon 2 hours treatment with 0.5 μg/ml RhoA activator II (RhoA-II) and Rac1 inhibition upon 1 hour treatment with 100 μM NSC23766 (NSC) on cortex mechanics in modMCF-7 (post-EMT) in suspended interphase MCF-7 cells (top row) and MCF-7 cells in mitotic arrest (bottom row). The last condition was compared to the first condition. (Post-EMT cells are referred to as modMCF-7 and mod-MCF-10A, respectively. Number of cells measured: Panels **c-e**: Interphase: MCF-7 n=36, esiRhoA n=36, esiRac1 n=35, modMCF-7 n=31, esiRhoA n=26 and esiRac1 n=30, Mitosis: MCF-7 n=22, esiRhoA n=26, esiRac1 n=24, modMCF-7 n=28, esiRhoA n=28 and esiRac1 n=28. Panels **f-h**: Interphase: MCF-7 n=24, modMCF-7 n=24, NSC n=22, RhoA-II n=22 and NSC + RhoA-II n=24. Mitosis MCF-7 n=24, modMCF-7 n=28, NSC n=22, RhoA-II n=24 and NSC + RhoA-II n=22. Measurements are representative for at least two independent experiments. ^ns^p> 0.05, *p < 0.05, **p < 0.01, ***p < 0.001).

### Rac1 and RhoA activity affect cortex mechanics in a cell-cycle-dependent manner

Since we observed a significant change in the activity of RhoA and Rac1 upon EMT, we asked how Rac1 and RhoA influence the mechanical properties of the cortex. For this purpose, we performed a knock-down of RhoA or Rac1 by RNA interference in MCF-7 cells which led to a decrease of the respective protein activity (Figure S4c-d). We then performed cell-mechanical measurements in interphase and mitosis.

We found that in interphase, cortical tension and elastic modulus significantly increased after Rac1 knock-down, while the phase shift did not show significant changes (Figure 4c-e, top row). By contrast, we observed a reduction of cortical tension and cortical stiffness as well as an increase in phase shift in mitotic cells. A decrease of mitotic cortical tension upon Rac1 knock-down was shown before^40^. We repeated this measurement for post-EMT cells and found that changes were qualitatively equal but stronger corresponding to the above-mentioned increased Rac1 activity in post-EMT conditions (Figure 4c-e). Furthermore, inhibition of Rac1 by NSC23766 (Figure S4c,d), a Rac1-GEF inhibitor^41^, showed a similar trend as Rac1 knock-down, for both interphase and mitosis (Figure 4f-h). In summary, Rac1 knockdown or inhibition made the cortex more contractile in interphase and less contractile, less stiff and more liquid-like in mitosis. Therefore, Rac1 knock-down or inhibition-induced changes are in all six cortex-mechanical parameters (three parameters for interphase and mitosis, respectively) opposite to EMT-induced changes of cortical mechanics. Accordingly, Rac1 knock-down or inhibition reverses the cortex-mechanical EMT phenotype (Figure 4c-e, last condition, Figure 4f-h, 3^rd^ condition).

In interphase and mitosis, cortical stiffness and cortical tension decreased after RhoA knock-down as expected from previous work (Figure 4c-e, bottom row)^6,7,12,40^. The phase shift increase in mitosis after RhoA knockdown indicates a fluidization of the mitotic cortex. Post-EMT, RhoA knockdown cells showed the same trend in cortex mechanical changes in mitosis albeit less strongly, and no significant changes in interphase, which is in accordance with our findings of EMT-induced reduction of RhoA activity. Furthermore, pharmacological RhoA activation by RhoA activator-II in post-EMT condition (Figure S4c,d), blocking its intrinsic and GAP-stimulated GTPase activity^42^, resulted in a trend opposite to RhoA knock-down with increased cortical stiffness and cortical tension (Figure 4f-h).

In summary, both, Rac1 and RhoA signaling caused cortex-mechanical changes. In the case of Rac1, changes of cortex stiffness and contractility were opposite in interphase and mitosis.

We would like to point out that the observed phenotype of RhoA knock-down in mitosis was opposite to the EMT-induced phenotype in spite of EMT-associated RhoA activity decrease. We conjecture that this difference is due to the additional activation of Rac1 during EMT. Accordingly, we found that the combined activation of RhoA and inhibition of Rac1 can rescue the cell mechanical phenotype in post-EMT cells as depicted in Figure 4f-h, last condition (NSC + RhoA-II).

### Mitotic rounding in confined spheroids is enhanced through EMT

So far, we studied mitotic cells in 2D culture, where mitotic cells can easily become round since there are no mechanical obstacles to rounding. Accordingly, in 2D culture, mitotic cells are round in pre and post-EMT conditions. However, under physiologically relevant conditions, cells are mechanically confined by other cells or stroma. We therefore asked whether EMT and corresponding increase of cortical tension indeed lead to enhanced mitotic rounding when cells are cultured in a mechanically confining environment. To this end, we grew breast epithelial cells in 3D culture embedded in non-degradable PEG Heparin gel. Over time, cells grew into spheroidal cell aggregates which were mechanically confined by the gel. We quantified the roundness of cells in mitosis before and after EMT induction after seven days of growth (MCF-7: Figure 5a-c, MCF-10A: Figure S5a). 12-15 hours before fixation, S-trityl-L-cysteine (2 *μ*M) was added to arrest mitotic cells in mitosis in order to enrich mitotic cells. Spheroids were imaged through confocal z-stacks. For each mitotic cell, the largest cross-sectional area of the cell was used to quantify roundness (Figure 5b, right panel, Materials and Methods). Furthermore, spheroid size was quantified by the largest cross-sectional area of the spheroid and the corresponding number of cells in this cross section. We found that increasing gel stiffness indeed reduces mitotic roundness and spheroid size most likely due to mechanical hindrance (Figure 5c,f, S5g). Furthermore, our data show an increase in mitotic roundness and spheroid size after EMT (MCF-7: Figure 5c,f, S5g, MCF10A: Figure S5e). In particular, for MCF-7, EMT-induced proliferation increase is stronger in spheroids than in 2D culture (Figure S5g and Figure S1d). For MCF-10A, proliferation trends through EMT are reversed in mechanical confinement: while EMT-induction reduces proliferation in 2D, proliferation is enhanced in spheroids (Figure S1d and S5e). These findings suggest that EMT-induced cell-mechanical changes enhance mitotic rounding and, thus, support successful and timely cell division in mechanically confining environments.

**Figure 5:**
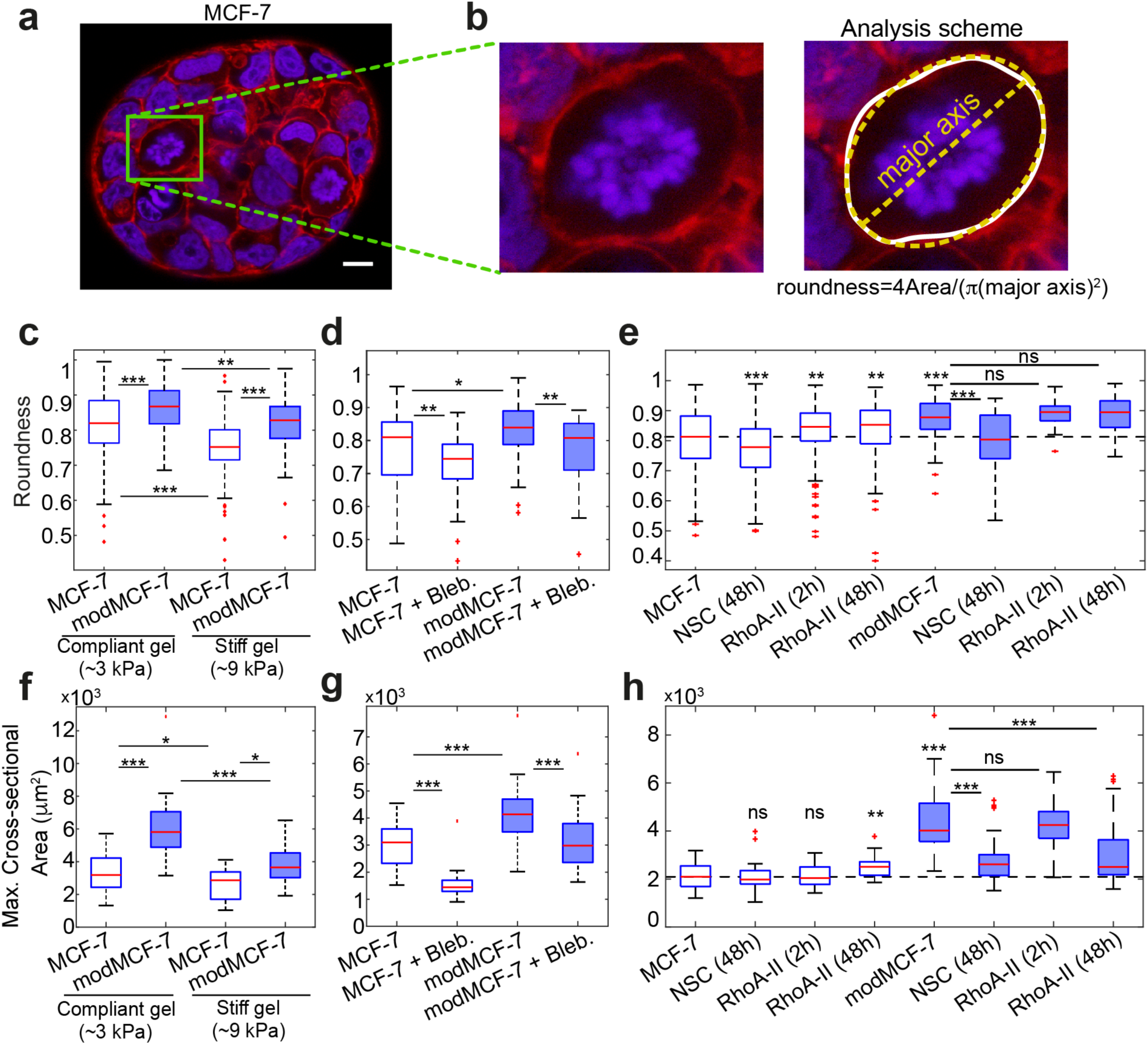
The influence of EMT on mitotic roundness in MCF-7 tumor spheroids. **a**, Confocal images of cross-sections of MCF-7 spheroids in PEG heparin gel after one week of growth. Cells were fixed and stained for DNA (DAPI, blue) and F-actin (phalloidin, red). Mitotic cells were arrested in mitosis through co-incubation with STC (2 μM) prior to fixation and exhibit condensed chromosomes (see e.g. cells in green frame). Scale bar: 10 μm. **b**, Left panel: zoom of cross-sections of mitotic cells in **a**. Right panel: For image analysis, the cell outline of the largest cell cross-section was first determined (white). Roundness of the cell was then calculated using Fiji including the fit of an ellipse (yellow), see Materials and Methods. **c**, Roundness of mitotic cells in spheroids before and after EMT in compliant and stiff hydrogels (Young’s modulus of ∼3 or ∼9 kPa, respectively). **d**, Roundness of mitotic cells in compliant gels with or without inhibition of cortical contractility through the myosin inhibitor blebbistatin (10 μM). **e**, Roundness of mitotic cells in compliant gels with or without Rac1 inhibition upon 48 hours treatment with 50 μM NSC23766 (NSC) and RhoA activation upon 2 hours or 48 hours treatment with 1 μg/ml RhoA activator-II (RhoA-II). **f**, Influence of EMT on spheroid size in compliant and stiff hydrogels. Spheroid size was quantified by the largest cross-sectional area of the spheroid. **g**, Influence of Myosin inhibition on spheroid size in compliant gels (Blebbistatin treatment for 2-days at 10 μM, see Materials and Methods) in pre- and post-EMT conditions. **h**, Influence of Rac1 inhibition on spheroid size upon 48 hours treatment with 50 μM NSC23766 (NSC) and RhoA activation upon 2 hours or 48 hours treatment with 1 μg/ml RhoA activator-II (RhoA-II) on spheroid size in pre- and post-EMT conditions. (Post-EMT cells are referred to as modMCF-7 and mod-MCF-10A, respectively. Number of mitotic cells analyzed: Panel **c**: Compliant gel, MCF-7 n=133, modMCF-7 n=92, Stiff gel, MCF-7 n=93 and modMCF-7 n=71. Panel **d**: MCF-7 n=26, MCF-7+Bleb., n=54, modMCF-7 n=58 and modMCF-7+Bleb. n=47. Panel **e**: MCF-7 n=155, NSC (48h) n=192, RhoA-II (2h) n=202, RhoA-II (48h) n=167, modMCF-7 n=71, NSC (48h) n=35, RhoA-II (2h) n=50 and RhoA-II (48h) n=83. Number of spheroids analyzed: Panel **f**: MCF-7 n=37, modMCF-7 n=34, (Stiff gel) MCF-7 n=17 and modMCF-7 n=15. Panel **g**: MCF-7= 26, MCF-7+Bleb. n=25, modMCF-7 n=31, modMCF-7+Bleb. n=33. Panel **h**: MCF-7 n=40, NSC (48h) n=40, RhoA-II (2h) n=40, RhoA-II (48h) n=31, modMCF-7 n=41, NSC (48h) n=39, RhoA-II (2h) n=40 and RhoA-II (48h) n=42. Measurements are representative for at least two independent experiments. ^ns^p> 0.05, *p < 0.05, **p < 0.01, ***p < 0.001).

**Figure 6:**
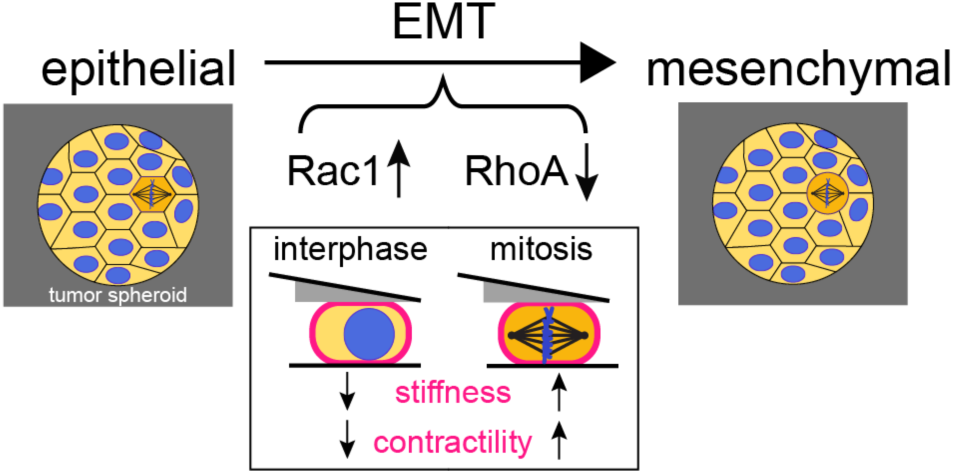
Our model on mechanical and molecular changes upon EMT: EMT in breast epithelial cells MCF-7 and MCF-10A leads to cortex-mechanical changes; cortical stiffness and contractility are reduced in interphase cells but increased in mitotic cells. Cell-mechanical changes upon EMT are accompanied by a strong increase in Rac1 activity and decrease in RhoA activity.

To identify whether cortical contractility contributed to this effect, we tested the effect of the myosin-inhibiting drug Blebbistatin on spheroids. Myosin inhibition reduces actomyosin contractility. Indeed, we find that Blebbistatin decreases mitotic rounding in spheroids and spheroid growth in pre- and post-EMT conditions (MCF-7: Figure 5d,g, S5h, MCF-10A: Figure S5a). Furthermore, our data indicate that spheroid growth enhancement of EMT-inducing TPA-treatment can be annulled through addition of Blebbistatin (MCF-7: Figure 4g, MCF-10A: Figure S5c,e). These findings show evidence that actomyosin contractility has an influence on mitotic rounding and cell proliferation in 3D culture.

To observe the influence of Rac1 and RhoA activity changes in 3D culture, we used again the Rac1-GEF inhibitor NSC23766 and RhoA activator-II; we show that these drugs induce corresponding changes in Rac1 and RhoA activity (MCF-7: Figure S4c,d). Rac1 inhibition resulted in reduction of roundness particularly in post-EMT conditions (MCF-7: Figure 5e, MCF-10A: Figure S5b). This reduction of roundness was accompanied by a significant reduction of spheroid size post-EMT (MCF-7: Figure 5h, S5i, MCF-10A: Figure S5d,f).

In pre-EMT MCF-7 spheroids, RhoA activation led to an increase of mitotic roundness and spheroid size (Figure 5e,h, S5i). By comparison, MCF-10A spheroids showed no significant changes in roundness and spheroid size (Figure S5,b,d,f).

In post-EMT spheroids, RhoA activation did not change mitotic roundness (MCF-7: Figure 5e, MCF-10A: Figure S5b). To our surprise, though, MCF-7 cell proliferation was reduced upon this treatment (Figure 5e,h, S5i). This proliferation reduction can thus not be related to altered mitotic rounding and needs to be further investigated in future research. In MCF-10A, RhoA activation had no effect on spheroid size.

## Discussion

Here, we have shown that pharmacologically-induced EMT gives rise to changes in mechanical properties and composition of the actin cell cortex of breast epithelial cells. In our study, we compared interphase cells in suspension and mitotic cells in 2D culture before and after EMT. In particular, we observed a softer, less contractile cortex in post-EMT interphase cells and a more contractile, stiffer cortex phenotype in post-EMT mitotic cells in comparison to the respective pre-EMT phenotype (Figure 2e-g). Concomitantly, we found corresponding changes of cortical myosin upon EMT: cortical myosin decreases in interphase cells but increases in mitotic cells (Figure 3a,b). These findings suggest that observed changes of cortical contractility upon EMT are, at least partly, caused by respective changes of myosin at the cortex. Furthermore, corresponding stress-stiffening or stress-stiffening release may account for stiffness changes in mitosis and interphase. Stress-stiffening in response to active stresses generated by myosin was reported in *in vitro* reconstituted actin networks and in cells^34,43–45^. It can be accounted for by different physical mechanisms^46–48^. In addition, we report an EMT-induced increase of cortical actin in mitotic cells which may further assist cortical stiffening and contractility due to associated structural changes of the cortical network (Figure 3a,b)^49–51^. Our findings put forward that the observed changes in actin and myosin are caused by upstream activity changes of Rho GTPases such as Rac1 and RhoA. The details of responsible signalling pathways will be the object of future research.

Our observation that post-EMT interphase cells have a softer actin cytoskeleton is in agreement with a study by Osborne *et al.* that reported softening of adherent cells upon EMT^52^ and with reports that metastatic cancer cells are on average softer than healthy and non-metastatic cells^20,53–55^. Here, we show for the first time opposite changes in cell cortex mechanics in interphase and mitotic cells upon EMT (Figure 2e-g).

Enhanced cortical contractility in mitosis corresponds to an enhanced cell surface tension in mitosis and provides an important mechanical driving force for cell rounding^1,3,5–11^. Therefore, our findings suggest increased mitotic rounding strength for post-EMT breast epithelial cells and thus an increased potency to undergo successful mitotic rounding and cell division in mechanical confinement. Furthermore, our observation of concurrent softening of interphase cells upon EMT may facilitate mitotic rounding further as post-EMT interphase cells in the neighborhood of a mitotic cell are more easily deformable. In fact, we could confirm enhanced mitotic rounding in post-EMT conditions by our observation of increased roundness of mitotic cells in spheroids growing in non-degradable Peg-Heparin hydrogels (Figure 5c). Furthermore, we observed increased growth of post-EMT MCF-7 and MCF-10A spheroids in the presence of intact actomyosin contractility (Figure 5f,g, S5a,c,e,g).

In search of a molecular explanation for the altered cortex mechanics, we found that activity of the Rho GTPase Rac1 is strongly increased in MCF-7 and MCF-10A cells after EMT which is in accordance with previous findings^56–60^ (Figure 4). In addition, we showed that Rac1 activity is a prerequisite of increased mitotic rounding and proliferation in post-EMT spheroids (Figure 5e,h, S5b,d,f,i). By contrast, we found a strongly decreased activity of the Rho GTPase RhoA after EMT (Figure 4a,b).

Our cell-mechanical study suggests that EMT-induced mechanical changes of the cortex could mainly be accounted for by the mechanical impact of EMT-induced Rac1 activation (Figure 4c-h); we show that decrease of Rac1 activity through gene knock-down and pharmacological inhibition induces cortex-mechanical changes that are opposite to EMT-induced changes in all six characterizing mechanical parameters (Figure 4c-h). Correspondingly, Rac1 inhibition rescues or overcompensates EMT-induced cell-mechanical changes in post-EMT cells.

In summary, we report here a cell-mechanical study that shows significant changes of cortex mechanics upon EMT and implies increased mitotic rounding strength of post-EMT breast epithelial cells. While EMT is commonly regarded to be connected to cancer progression through the promotion of cancer cell invasiveness, our findings indicate that EMT might also support mitotic cell rounding and preceding successful cell division in the increasingly crowded environment of a growing tumor.

## Materials and Methods

### Cell culture

The cultured cells were maintained as follows: MCF-7 cells were grown in RPMI-1640 medium (PN:2187-034, life technologies) supplemented with 10% (v/v) fetal bovine serum, 100 *μ*g/ml penicillin, 100 *μ*g/ml streptomycin (all Invitrogen) at 37°Cwith 5% CO_2_. MCF-10A cells were cultured in DMEM/F12 medium (PN:11330-032, Invitrogen) supplemented with 5% (v/v) horse serum (PN:16050-122, Invitrogen), 100 *μ*g/ml penicillin, 100 *μ*g/ml streptomycin (all Invitrogen), 20 mg/ml epidermal growth factor (PN:AF-100-15, Peprotech), 0.5 mg/ml hydrocortisone (PN:H-0888, Sigma), 100 ng/ml cholera toxin (PN:C-8052, Sigma), 10 *μ*g/ml insulin (PN:I-1882, Sigma) at 37°Cwith 5% CO_2_.

In MCF-7 cells, EMT was induced by incubating cells in medium supplemented with 100 nM 12-*O*-tetradecanoylphorbol-13-acetate (TPA) (PN:P8139, Sigma) for 48 hours prior to measurement ^26^. In MCF-10A, EMT was induced by incubating cells in medium supplemented with 10 ng/ml TGF-*β*1 (PN:90900-1, BPSBioscience, San Diego, USA) one week prior to measurement^27–29^. Measured cell volumes before and after EMT induction are depicted in Figure S2. For MCF-10A cells, we observe an average increase of cell volumes in mitosis of ∼15%.

Rac1 inhibition was performed by treatment with NSC23766 (PN:2161, Tocris). RhoA activation was achieved by treatment with RhoA activator-II (PN:CN03, Cytoskeleton, Inc.).

### AFM measurements of cells

To prepare mitotic cells for AFM measurements, we seeded ∼10.000 cells in a silicon cultivation chamber (0.56 cm^2^ area, from ibidi 12-well chamber) that was placed in a 35 mm cell culture dish (fluorodish FD35-100, glass bottom, World Precision Instruments) one day before the measurement so that a confluency of approximately 30% was reached on the measurement day. We induced mitotic arrest by supplementing S-trityl-L-cysteine (STC, Sigma) 2-8 hours before the measurement at a concentration of 2 *μ*M. For measurement, mitotically arrested cells were identified by their shape. Their uncompressed diameter ranged typically from 18-23 *μ*m. AFM measurement of metaphase cells (without addition of STC) showed similar measured mechanical parameters and trend upon Rac1 and RhoA knock-down in comparison to STC-treated cells (Figure S6) indicating that STC does not significantly interfere with GTPase signaling of our measurements.

To prepare AFM measurements of suspended interphase cells, cell culture dishes (fluorodish FD35-100) were coated by incubating the dish at 37°Cwith 0.5 mg/ml PLL-g-PEG dissolved in PBS for 1 hour (SuSoS, Dubendorf, Switzerland) to prevent cell adhesion. Prior to measurements, cultured cells were detached by the addition of 1X 0.05% trypsin-EDTA (Invitrogen). Approximately 30.000 cells in suspension were placed in the coated culture dish. The culture medium was changed to CO_2_-independent DMEM (RN: 12800-017, Invitrogen) with 4 mM NaHCO_3_ buffered with 20 mM HEPES/NaOH pH 7.2, for AFM experiments approximately 2 hours before the measurement^8,9,24^.

The experimental set-up included an AFM (Nanowizard I, JPK Instruments) that was mounted on a Zeiss Axiovert 200M optical, wide-field microscope. 20x objective (Zeiss, Plan Apochromat, NA=0.8) along with a CCD camera (DMK 23U445 from Theimagingsource). Cell culture dishes were kept in a petri dish heater (JPK Instruments) at 37°Cduring the experiment. Prior to every experiment, the spring constant of the cantilever was calibrated by thermal noise analysis (built-in software, JPK) using a correction factor of 0.817 for rectangular cantilevers^61^. The cantilevers used were tipless, 200 – 350 *μ*m long, 35 *μ*m wide and 2 *μ*m thick (NSC12/tipless/no Aluminium or CSC37/tipless/no Aluminium, Mikromasch). The nominal force constants of the cantilevers ranged between 0.2 – 0.8 N/m. The cantilevers were supplied with a wedge, consisting of UV curing adhesive (Norland 63, Norland Products Inc.) to correct for the 10° tilt^62^. The measured force, piezo height and time were output at a time resolution of 500 Hz.

#### Oscillatory cell compression protocol

Preceding every cell compression, the AFM cantilever was lowered to the dish bottom in the vicinity of the cell until it touched the surface and then retracted to approximately 15 *μ*m above the surface. Subsequently, the free cantilever was moved and placed on top of the cell. Thereupon, a bright-field image of the equatorial plane of the confined cell is recorded in order to evaluate the equatorial cross-sectional area at a defined cell height, which in turn is used to estimate the cell volume as described in earlier work^24^. As the cell is sandwiched between the dish bottom and cantilever wedge, oscillatory height modulations of the AFM cantilever were undertaken, with piezo oscillation amplitudes of 0.25 *μ*m at a frequency of 1 Hz if not indicated otherwise. Throughout this step, the cell was typically maintained at a height corresponding to 60-70% of the original cell height. Using this degree of confinement, we verified that the major contribution to cell-mechanical response was the actin cortex and not the nucleus or the cytoplasm of the cell. This was done by comparative measurements with and without cortex-affecting cytoskeletal drugs (Latrunculin A and Blebbistatin), (Figure S7). Oscillatory cantilever height modulations were produced by a piecewise linear approximation of the sine-function^24^. The force acting on the cantilever and the height of the cantilever was constantly recorded. The height of the confined cell was calculated as the difference between the height that the cantilever was elevated above the dish surface and lowered onto the cell with the addition of the spike heights at the boundary of the wedge (formed by imperfections during the manufacturing process^62^) and the force-induced deflection of the cantilever. We estimate a total error of the cell height of approximately 0.5 *μ*m due to irregularities of the cantilever wedge and vertical movement of the cantilever to a position above the cell.

#### Data analysis of AFM measurements

The data analysis procedure has been described in detail in earlier work^24^. In brief, the time series of measured AFM force *F* and AFM cantilever height *h* were analyzed in the following way: Data analysis of oscillatory measurements were performed after a transient relaxation phase in a time interval of steady state oscillations (constant force and constant cantilever height oscillation amplitude, see Figure 2). In an analysis time window of 5-10 periods, we estimated for every time point effective tension *γ*_*eff*_ = *F*_*AFM*_/*A*_*con*_(1/*R*_1_ + 1/*R*_2_) and relative cell surface area change (area strain) *ε*_*A*_=(*A* − ⟨*A*⟩)/⟨*A*⟩. Here *R*_1_, *R*_2_ and *A*_*con*_ denote geometrical parameters of cell shape (radii of curvature and contact area), while *A* indicates the cell surface area and ⟨*A*⟩ denotes its time-averaged value. Geometrical parameters of cell shape are calculated assuming a shape of minimal surface area at constant cell volume as described before^34^. Cortical tension *γ* was estimated as the time-averaged value of *γ*_*eff*_ during an oscillation period.

For a rheological analysis, effective cortical tension *γ*_*eff*_ is interpreted as stress readout, while *ε*_*A*_ is interpreted as strain readout. An amplitude and a phase angle associated to the oscillatory time variation of stress and strain are extracted. To estimate the value of the complex elastic modulus at a distinct frequency, we determine the phase angles *φ*_*γ*_ and 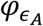 as well as amplitudes 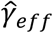 and 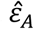 of effective tension and surface area strain, respectively. Cortical stiffness was calculated as 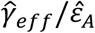 while the phase shift was determined as *φ* = (*φ*_*γ*_ − *φ*_*A*_). Statistical analysis of cortex mechanical parameters was performed in Matlab using the commands ‘boxplot’ and ‘ranksum’ to generate boxplots and determine p-values from a Mann-Whitney U-test (two-tailed), respectively.

### Plasmids and transfection

Cells were transiently transfected with plasmid DNA using Lipofectamine 2000 (PN: 11668030, Invitrogen), according to the manufacturer’s protocol. To achieve post-EMT conditions, MCF-7 cells were seeded and treated with TPA one day before transfection. Cells were imaged one day following transfection. The plasmid MApple-LC-Myosin-N-7 was a gift from Michael Davidson (Addgene plasmid # 54920; http://n2t.net/addgene:54920; RRID:Addgene_54920). The plasmid MCherry-Actin-C-18 was a gift from Michael Davidson (Addgene plasmid # 54967; http://n2t.net/addgene:54967; RRID:Addgene_54967).

### Confocal imaging of transfected cells

The cells were transferred to PLL-g-PEG coated fluorodishes (FD35-100) with CO_2_-independent culture medium (described previously). Cells were stained with Hoechst 33342 solution (PN:62249, Invitrogen) in order to distinguish between mitotic and interphase cells. During imaging they were maintained at 37°Cusing ibidi heating stage. Images were recorded using a Zeiss LSM700 confocal microscope of the CMCB light microscopy facility, incorporating a Zeiss C-Apochromat 40x/1.2 water objective. Images were taken at the equatorial diameter of each cell exhibiting the largest cross-sectional area.

### Quantification cortex-associated actin and myosin from confocal imaging

For image analysis of confocal images, the cell boundary was determined by matlab custom-code (Figure S3b shows example cell, the cell boundary is marked in red). Along this cell boundary 200 radial, equidistant lines were determined extending 2 *μ*m to the cell interior and 2 *μ*m into the exterior (Figure S3b, red lines orthogonal to cell boundary, only every 10^th^ line is plotted out of 200). The radial fluorescence profiles corresponding to these lines were averaged over all 200 lines (Figure S3c, blue curve). This averaged intensity profile is then fitted by the function:

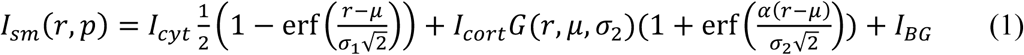

with fit parameters *p* = {*μ, σ*_1_, *σ*_2_, *I*_*cyt*_, *I*_*cort*_, *I*_*BG*_, *α*}. In this way, we obtain a smoothed variant *I*_*sm*_(*r, p*) of the intensity profile along the radial coordinate. A respective exemplary fit of *I*_*sm*_(*r, p*) is shown in Figure S3c, orange curve. Here, erf(*r*) denotes the error function and G(*r, μ, σ*_2_) denotes a Gaussian function with mean value *μ* and standard deviation *σ*_2_. The term 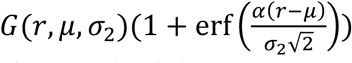 is a corresponding skewed Gaussian function with skewness characterized by parameter *α*. The parameters *I*_*cyt*_ and *I*_*cort*_ quantify the amplitude of cytoplasmic versus cortical fluorescence, while *σ*_1_ and *σ*_2_ quantify the smearing out of the cytoplasmic or cortical fluorescence distribution, respectively. *I*_*BG*_ captures the value of background intensity, while *μ* is the radial coordinate of cell boundary localization.

The skewed Gaussian contribution in the fitted intensity profile (Figure S3c, orange curve) is interpreted as fluorescence contribution from the cortical proteins, while the remaining part is interpreted as a contribution of cytoplasmic fluorescence and background fluorescence *I*_*BG*_. The skewed Gaussian was then integrated along the radial direction to obtain *I*_*2D,cortex*_ as a measure for the 2D concentration of fluorescent protein at the cortex. This integrated intensity was then normalised by the intensity *I*_*cyt*_ of the cytoplasm. The normalised intensity *I*_*2D,cortex*_ /*I*_*cyt*_ is then used as a measure of cortex-to-cytoplasm ratio in boxplots in Figure 3b.

### Immunostaining and confocal imaging of Phalloidin-labelled cells

The suspended (interphase or STC-arrested mitotic) cells were fixed with 4% formaldehyde/PBS for 1 h, followed by a 30 min permeabilization step in 0.2% Triton X-100. The cells were then stained for 1 h with 5 µg/ml DAPI and 0.2µg/ml Phalloidin-Alexa Fluor 647 in 2% BSA/PBS solution. Images were recorded using a Zeiss LSM700 confocal microscope of the CMCB light microscopy facility, incorporating a Zeiss C-Apochromat 40x/1.2 water objective. Images were taken at the equatorial diameter of each cell exhibiting the largest cross-sectional area.

### Estimation of cortical thickness

Prior to the measurement, fluorescent labels of F-actin and of the plasma membrane were introduced into MCF-7 cells following the respective protocols of the manufacturer: we transfected cells with a plasmid expressing Lifeact-GFP two days prior to the measurement using Lipofectamine 2000; furthermore, we employed the membrane dye CellMask™ Orange (PN: C10045, Thermofisher). Mitotic or non-adherent round interphase cells were imaged in two fluorescence channels (GFP and mOrange) with an LSM700 confocal microscope using a C-Apochromat 40x water-immersion objective with an NA of 1.2. We recorded z-stacks with an interval of 200 or 250 nm and a total of 25 slices. Afterwards, the slice with the largest cross-sectional area was chosen in Fiji. This slice was afterwards analysed with Matlab custom-code. For both channels, the cell boundary was determined and a lateral shift between the two channels was removed by calculation of the difference of the respective centres of gravity. Then, fluorescence of either channel was averaged along the cortical circumference and Equation 1 was fit to the radial fluorescence profiles with a skewness value of zero following the description of ^63^ (see Figure S3c for an exemplary plot). The radial coordinate *μ* of the peak localization of the fitted Gaussian functions was output for fluorescence of CellMask™ Orange and Lifeact-GFP. The twofold difference between these peak localizations was taken as a cortex thickness estimate as has been described before^63^ (Figure S3c). Chromatic shifts of the z-position of the focal plane were estimated to be smaller than 400 nm. Associated errors to our cortex thickness measurements were estimated to be less than 10% given typical equatorial cell diameters between 15 and 20 *μ*m.

### FRAP measurements at the cortex

To monitor turnover of the actin cortex, we transfected MCF-7 cells with a plasmid expressing mCherry-ActB as described above. During measurements, mitotic or non-adherent interphase cells were imaged in a time series with a confocal setup (Zeiss LSM 700, 20x objective Plan Apochromat, NA=0.8). At the beginning of the time series, a small region of the cortex was photobleached in the equatorial plane of the round cell. Afterwards, cell fluorescence in the equatorial plane was recorded for a total time span of 120 s at a time interval of 3 s (Figure S3g). For image analysis, the cell circumference and the bleached region of the cortex were determined by image analysis with Matlab custom-code as described in^64^. Averaged fluorescence in the bleached region was output as a function of time. Where necessary, translational motion of cells during FRAP recovery was removed from the movie by the StackReg plugin of Fiji before further analysis.

### Western blotting

Protein expression in MCF-10A and MCF-7 before and after EMT was analyzed using Western blotting. Cells were seeded onto a 6-well plate and grown up to a confluency of 80-90% with or without EMT-inducing agents. Thereafter, cells were lysed in SDS sample/lysis buffer (62.5 mM TrisHcl pH 6.8, 2% SDS, 10% Glycerol, 50 mM DTT and 0.01% Bromophenolblue).

For analysis of protein expression in mitotic cells, STC was added to a concentration of 2 *μ*M to the cell medium 12 h before cells were harvested in order to enrich mitotic cells. For harvesting, mitotic cells were collected by shake-off.

Cell lysates were incubated for 1 hour with the lysis buffer at 4°C. They were then centrifuged at 15,000 rpm for 5 min at 4°C to discard insoluble content and then boiled for 5 minutes. 15 *μ*l of the cleared lysate was then used for immunoblotting. The cleared lysates were first run on precast protein gels (PN:456-1096 or 456-1093, Bio-Rad) in MOPS SDS running buffer (B0001, Invitrogen). Subsequently, proteins were transferred to PVDF membranes (Millipore).

PVDF membranes were blocked with 5% (w/v) skimmed milk powder (T145.1, Carl Roth, Karlsruhe, Germany) in TBST (20 mM/L Tris-HCl, 137 mM/L NaCl, 0.1% Tween 20 (pH 7.6)) for 1 h at room temperature followed by washing with TBST, and incubation at 4°Covernight with the corresponding primary antibody diluted 1:1000 (Vimentin, E-Cadherin, N-Cadherin, Rac1) or 1:500 (RhoA, Cdc42) or 1:5000 (GAPDH) in 3% (w/v) bovine serum albumin/TBST solution. Thereupon, the blots were incubated with appropriate secondary antibodies conjugated to horseradish peroxidase, Rabbit anti-mouse HRP (PN:ab97045, Abcam) or Goat anti-rabbit HRP (PN:ab97051, Abcam) at 1:3000 dilution in 5% (w/v) skimmed milk powder in TBST for 1 h at room temperature. After TBST washings, specifically bound antibodies were detected using Pierce enhanced chemiluminescence substrate (ECL) (PN:32109, Invitrogen). The bands were visualized and analyzed using a CCD-based digital blot scanner, ImageQuant LAS4000 (GE Healthcare Europe, Freiburg, Germany).

Primary antibodies used are as follows: E-Cadherin (PN:5296, Cell Signaling Technology), Vimentin (PN:AMF-17b, Developmental Studies Hybridoma Bank, Iowa City, IA), N-Cadherin (PN:13116, Cell Signaling Technologies), GAPDH (PN:ab9485, Abcam), RhoA (PN:ARH04, Cytoskeleton Inc.), Cdc42 (PN:ACD03, Cytoskeleton Inc.), Rac1 (PN:PA1091X, Invitrogen), ACTA (PN:ab3280, Abcam) and MLC2 (PN:8505, Cell Signaling Technologies). P-values of Western blots were determined with Mathematica using a univariate t-test testing the hypothesis that the distribution of relative changes in control conditions is either larger or smaller than one.

### Active Rho-GTPase pull-down assay

Pull-down assays were performed for RhoA-GTP, Rac1-GTP and Cdc42-GTP using the RhoA and Cdc42 pull-down activation assay biochem kit (PN:BK036-S and BK034-S, Cytoskeleton, Inc.) following the protocol provided by the manufacturer. Briefly, cells were lysed in lysis buffer and supplemented with 1 mM Protease Inhibitor Cocktail. Cell lysates for pull-down were harvested rapidly on ice with a cell scraper.

For pull-down of active forms of Rho GTPases, a 50 *μ*l volume of cleared cell lysate was added to a pre-determined amount of rhotekin-RBD beads (for RhoA) or PAK-PBD beads (for Rac1 and Cdc42) at 4°C. Lysate with beads was incubated for 60 min with gentle rocking. Then, the solution was centrifuged at 15,000 rpm for 1 min at 4°C and the supernatant was removed. The pellet (including the beads) was washed in washing buffer. Beads were then resuspended in 20 *μ*l of Laemmli sample buffer, boiled for 2 min.

As a positive and negative control, bead-aided pull-down was repeated with cell lysates previously mixed with non-hydrolysable GTPγS or GDP (Figure 4a, first and second column). Beads from pull-down and full lysate were subsequently analyzed in different lanes by SDS-PAGE and Western blot as described in the previous section.

### Gene knock-down through RNA interference

We performed gene knock-downs through RNA interference (RNAi). Cells were transfected with primary esiRNA (Eupheria Biotech, Dresden, Germany) targeting the genes Rac1 and RhoA at a concentration of 25 nM, using the transfection reagent Lipofectamin RNAiMax (Invitrogen). For a negative control, we used firefly luciferase esiRNA (F-Luc). For a positive control, we used Eg5/kif11 esiRNA, which in all cases resulted in mitotic arrest of more than 60% of the cells (indicating a transfection efficiency of more than 60% for each experiment). At day 0, 30,000 cells were seeded into a 24-well plate (NuncMicroWell Plates with Nunclon; Thermo Fisher Scientific, Waltham, MA). Transfection was carried out at day 1, medium was exchanged at day 2 (without antibiotics), and transfected cells were measured at day 3. For mitotic cells, approximately 12 hours before measurements, cells were detached, diluted, and transferred onto a glass-bottom petri dish (FD35-100, World Precision Instruments) to obtain a confluency of 20 – 40% during AFM measurements. For interphase cells, 2-3 hours before measurement the cells were detached and transferred to PLL-g-PEG-coated petri dishes (see Section on AFM measurements of cells).

Measured cell volumes before and after gene knock-downs are depicted in Figures S4, respectively. Our data indicate a decrease in cell volume upon Rac1 knock-down pre-EMT in mitotic cells and a reduction in interphase cell volume for post-EMT conditions (Figure S4e).

### Preparation of spheroids in starPEG-Heparin hydrogels

*In situ* assembling biohybrid multiarmed poly(ethylene glycol) (starPEG)-heparin hydrogels, prepared as previously reported^56^, were used for embedding cells to grow into spheroids. In brief, cells were detached from the tissue flasks and resuspended in PBS. Subsequently, they were mixed with freshly prepared heparin-maleimide-8 solution at a density of approximately 5 x 10^5^ cells/ml, and a final heparin-maleimide concentration of 3 mM^25^. PEG precursors were reconstituted at a concentration of 3 mM and put in an ultrasonic bath for 10-20s in order to dissolve. We then mixed 15 µl of the heparin-cell suspension with equal volume of PEG solution in a pre-chilled Eppendorf tube, using a low binding pipette tip. Thereafter, a 25 µl drop of PEG-heparin-cell mixture was pipetted onto a hydrophobic glass slide. After gelling, hydrogels were gently detached from the glass slide and placed in a 24-well plate supplemented with the cell culture medium. The spheroids were allowed to grow in the non-degradable hydrogel constructs for 7 days including the chemical treatment time for EMT-induction, before being fixed and immunostained. Medium was exchanged every second day. After seven days of growth, the medium was supplemented with 1 µM STC to enrich mitotic cells 13-15 h prior to fixation. Blebbistatin, NSC23766 and RhoA activator II were added to a concentration of 10 *μ*M, 50 *μ*M and 1 *μ*g/ml, respectively. Drugs were added either two hours prior to addition of STC to test its effect on mitotic rounding (data in Figure 5) or 48 hours prior to addition of STC to test its effect on cell proliferation and spheroid growth (data in Figure 5 and Figure S5).

### Quantification of hydrogel stiffness

Disc-shaped hydrogels, not seeded with cells, were immersed in PBS after preparation and kept at 4 °C to the following day for mechanical quantification. After equilibrating them at room temperature for 1 hour, gels were mounted onto glass object slides using Cell-Tak™ (Thermofisher). Gel stiffness was probed by AFM indentation^18,66,67^. Cantilevers (arrow T1, Nanoworld), that had been modified with polystyrene beads of 10 *μ*m diameter (Microparticles GmbH) using epoxy glue (Araldite), were calibrated using the thermal noise method implemented in the AFM software and probed at room temperature in PBS using a speed of 5 *μ*m/sec and a relative force setpoint of 2.5 nN. Force distance curves were processed using the JPK data processing software. Indentation parts of the force curves (about 1 *μ*m depth) were fitted using the Hertz model for a spherical indenter, assuming a Poisson ratio of 0.5.

### Immunostaining and confocal imaging of spheroids in hydrogel constructs

The spheroid hydrogel constructs were fixed at day 7 with 4% formaldehyde/PBS for 1 h, followed by a 30 min permeabilization step in 0.2% Triton X-100. The spheroids were then stained for 4 h with 5 µg/ml DAPI and 0.2µg/ml Phalloidin-Alexa Fluor 488 in 2% BSA/PBS solution. Then hydrogels were immersed at least for 1 hour in PBS. For imaging, hydrogels were cut horizontally with a blade and were placed on a cover slip to image the interior of the hydrogel. Per hydrogel 5-15 spheroids were imaged recording a confocal z-stack with a Zeiss LSM700 confocal microscope of the CMCB light microscopy facility, using a Zeiss C-Apochromat 40x/1.2 water objective (z-step: 0.4-0.9 µm).

### Image analysis of spheroid confocal images

The software Fiji was used to characterize morphology of mitotic cells inside spheroids. We identified mitotic cells inside a spheroid from spheroid z-stacks (red channel, F-actin, blue channel, DNA) by observing DNA structure. The largest cross-sectional area in the z-stack of each mitotic cell was determined manually by actin fluorescence, which was then used to calculate the cross-sectional shape factor roundness. Roundness is determined using Fiji by fitting an ellipse to the cross-section and calculating 4(*Area*)/(*π*(*major axis*)^2^), where *Area* is the cross-sectional area. Furthermore, spheroid size was quantified by the largest cross-sectional area of the spheroid (Figure 4f).

Statistical analysis of mitotic cell roundness in different conditions was performed in Matlab using the commands ‘boxplot’ and ‘ranksum’ to generate boxplots and determine p-values from a Mann-Whitney U-test (two-tailed), respectively.

### Code Availability

Custom codes written on Matlab can be obtained from the corresponding author on reasonable request.

### Data Availability

All data supporting the findings of this study are available from the corresponding author on reasonable request.

## Acknowledgements

We thank Jörg Mansfeld, Mark Leaver, Carsten Hoege and Benjamin Friedrich for critical reading of the manuscript and helpful comments and Alf Honigmann and Stephan Grill for fruitful discussions on the project. Furthermore, we thank Jochen Guck, Isabel Richter and Shada Abuhattum for access and introduction to infrastructure in the lab. In addition, we thank the CMCB light microscopy facility for excellent support, in particular Ali Gheisari. We also thank Marta Urbanska for the provision of Lifeact-GFP plasmid.

KH and EFF were supported by the Deutsche Forschungsgemeinschaft (DFG, German Research Foundation) under Germany’s Excellence Strategy – EXC-2068 – 390729961– Cluster of Excellence Physics of Life of TU Dresden.

## Author Contributions

K.H. performed the experiments. K.H. and E.F.F. designed the experiments and performed data analysis. K.H. and A.T. performed experiments on spheroids and provided reagents. C.W. provided reagents for hydrogel production in spheroid experiments. K.H. and E.F.F. wrote the manuscript.

