## Supplementary Information for "EMT-induced cell mechanical changes enhance mitotic rounding strength"

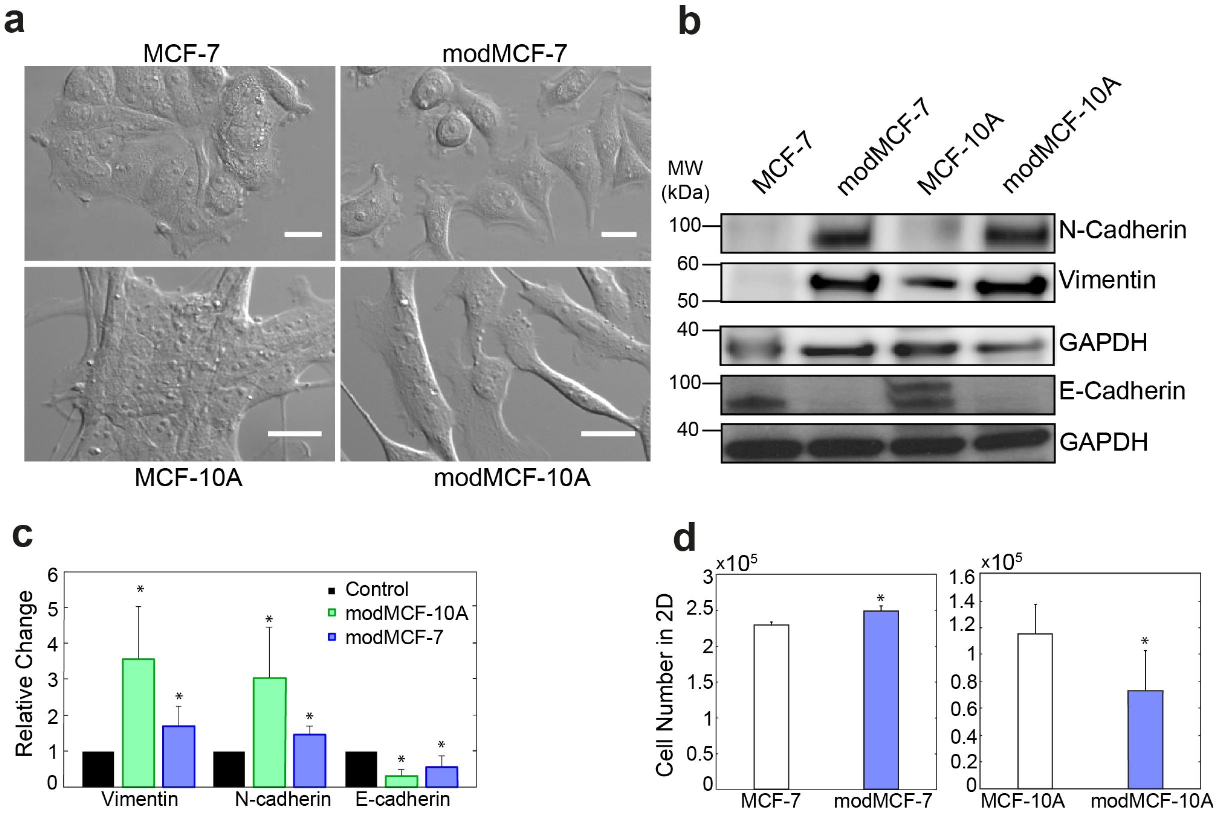


*Figure S1: Pharmacological induction of EMT in MCF-7 and MCF-10A cells through TPA and TGF-β1, respectively.* ***a,*** *Cell morphological changes upon EMT in MCF-7 (top row) and MCF-10A (bottom row): Untreated MCF-7 and MCF-10A cells grew in clusters while EMT-transformed cells grew more isolated from each other and show a trend towards a more spindle-shaped phenotype. Scale bar: top row: 10 µm, bottom row: 20 µm.* ***b****, Protein expression changes upon EMT: Western blots of E-cadherin (epithelial marker), N-cadherin and Vimentin (mesenchymal markers) from cell lysates before and after EMT. Post-EMT cells are referred to as modMCF-7 and mod-MCF-10A, respectively.* ***c****, Relative change of protein levels of epithelial marker E-cadherin and mesenchymal markers N-cadherin and Vimentin in breast epithelial cells before and after EMT from Western blot assays (E-Cadherin n=3, N-Cadherin n=3 and Vimentin n=4). Error bars indicate standard error of the mean.* ***d****, Number of cells in 2D upon 100 nM TPA treatment for 48 hours. P-values calculated with two-tailed student t-test. Error bars indicate standard deviations. Panel* ***d****: MCF-7 n=3, modMCF-7 n=3, MCF-10A n=3 and modMCF-10A n=3. (Post-EMT cells are referred to as modMCF-7 and mod-MCF-10A, respectively.* $^{ns}p > 0.05$, $*p < 0.05$, $**p < 0.01$, $***p < 0.001$*).*

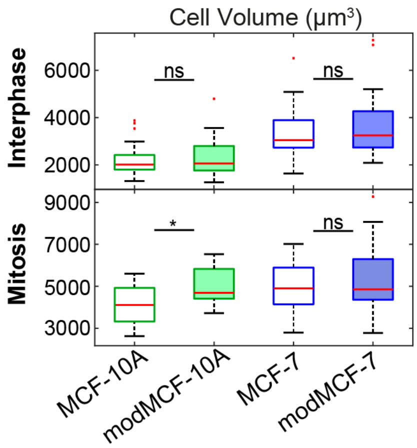


*Figure S2: Cell volumes of MCF-7 and MCF-10A cells before and after EMT corresponding to measurements presented in Fig. 2, main text, for suspended interphase cells (top) and cells in mitotic arrest (bottom) before and after EMT. (Post-EMT cells are referred to as modMCF-7 and mod-MCF-10A, respectively. Number of cells measured: Interphase: MCF-7 n=27, modMCF-7 n=28, MCF-10A n=27. Mitosis: MCF-7 n=34, modMCF-7 n=45, MCF-10A n=12, modMCF-10A n=12. $^{ns}p > 0.05$, $^{*}p < 0.05$, $^{**}p < 0.01$, $^{***}p < 0.001$).*

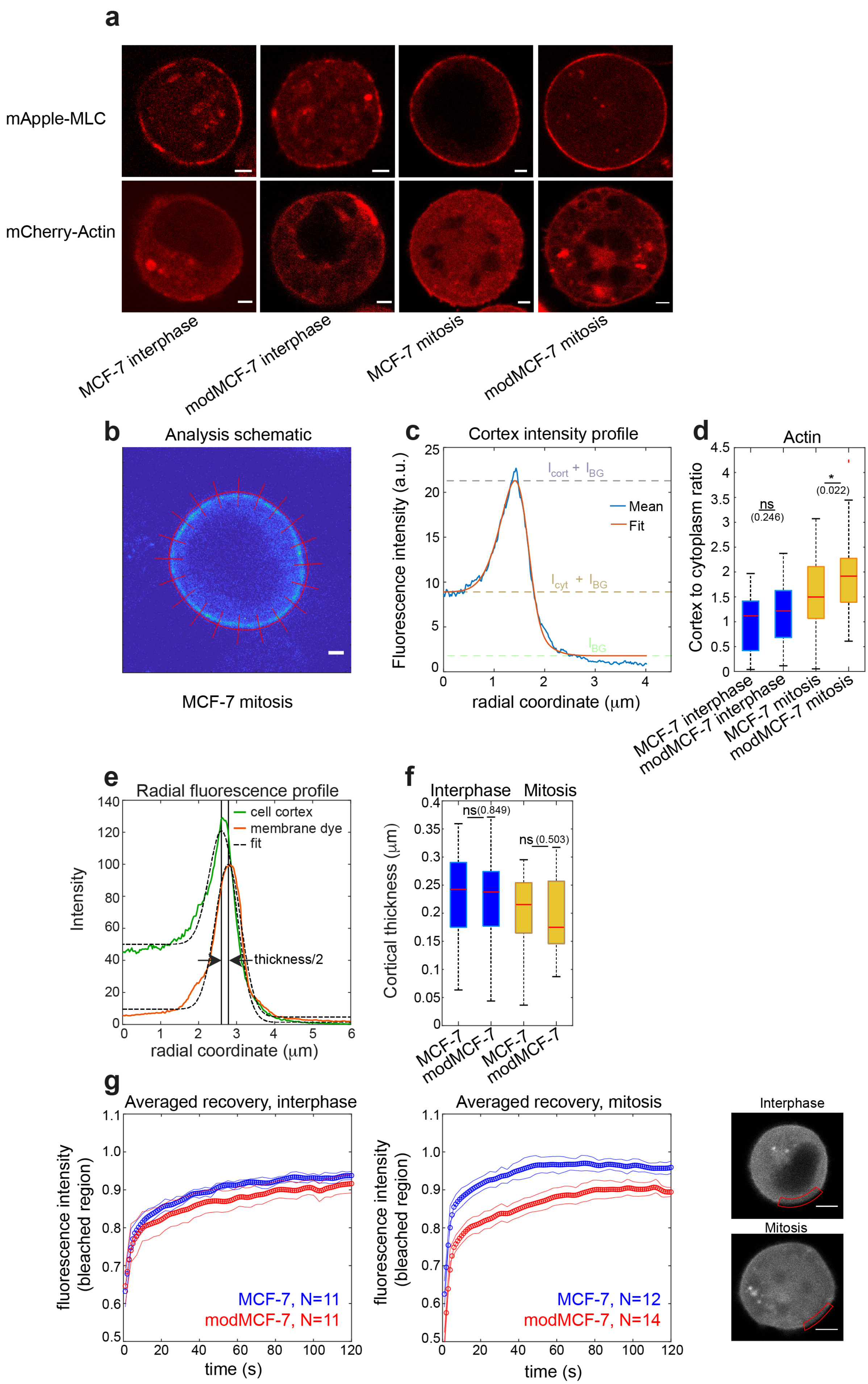
a
mApple-MLC
mCherry-Actin
MCF-7 interphase
modMCF-7 interphase
MCF-7 mitosis
modMCF-7 mitosis
b
Analysis schematic
MCF-7 mitosis
c
Cortex intensity profile
Fluorescence intensity (a.u.)
radial coordinate (μm)
Mean
Fit
d
Actin
Cortex to cytoplasm ratio
ns
(0.246)
*
(0.022)
e
Radial fluorescence profile
Intensity
cell cortex
membrane dye
fit
thickness/2
radial coordinate (μm)
f
Interphase
Mitosis
Cortical thickness (μm)
ns (0.849)
ns (0.503)
MCF-7
modMCF-7
g
Averaged recovery, interphase
Averaged recovery, mitosis
fluorescence intensity (bleached region)
MCF-7, N=11
modMCF-7, N=11
MCF-7, N=12
modMCF-7, N=14
time (s)
Interphase
Mitosis

*Figure S3: Actin and myosin changes in the cortex upon EMT.* ***a,*** *Representative confocal images of suspended interphase cells and STC-arrested mitotic cells expressing mCherry-ACTB or mApple-Myl9, pre- and post-EMT. Scale bar: 2 μm.* ***b,*** *Exemplary picture of myosin fluorescence profile of the equatorial cross-section of a mitotic cell including elements of image analysis. Scale bar: 2 μm.* ***c,*** *Average radial fluorescence intensity profile along red lines in* ***a*** *(blue curve). The fitted intensity profile, $I_{sm}(r,p)$, (see Materials and Methods, Equation 1) is shown in orange.* ***d,*** *Ratio of cortical versus cytoplasmic f-actin (labelled with phalloidin) in fixed suspended interphase cells and STC-arrested mitotic cells, pre- and post-EMT.* ***e,*** *Fluorescence intensity profiles along radial lines (see panel b) averaged along the equatorial cell circumference for Lifeact-GFP (green curve) and orange-stained membrane (orange curve). Fitted intensity profiles (see Materials and Methods) are shown in black.* ***f,*** *Cortical thickness estimate of suspended interphase and STC-arrested mitotic cells, pre- and post-EMT.* ***g,*** *Averaged recovery curves of normalized fluorescence intensity MCF-7 (blue curve) and modMCF-7 (red curve) in interphase (left panel) and mitosis (middle panel). Right panel shows exemplary confocal image of the equatorial plane of an interphase (top) and mitotic (bottom) cell with the bleached cortical region highlighted in red. Scale bar: 5 μm. (Post- EMT cells are referred to as modMCF-7. Number of cells measured: Panel* ***d****: Actin: MCF-7 interphase n=35, modMCF-7 interphase n=38, MCF-7 mitotic n=32, modMCF-7 mitotic n=36. Panel* ***f:*** *MCF-7 interphase n=20, modMCF-7 interphase n=18, MCF-7 mitotic n=23 and modMCF-7 mitotic n=23,. Panel* ***g:*** *MCF-7 interphase n=11, modMCF-7 interphase n=11, MCF-7 mitotic n=12, modMCF-7 mitotic n=14. $^{ns}p > 0.05$, $^{*}p < 0.05$, $^{**}p < 0.01$, $^{***}p < 0.001$).*

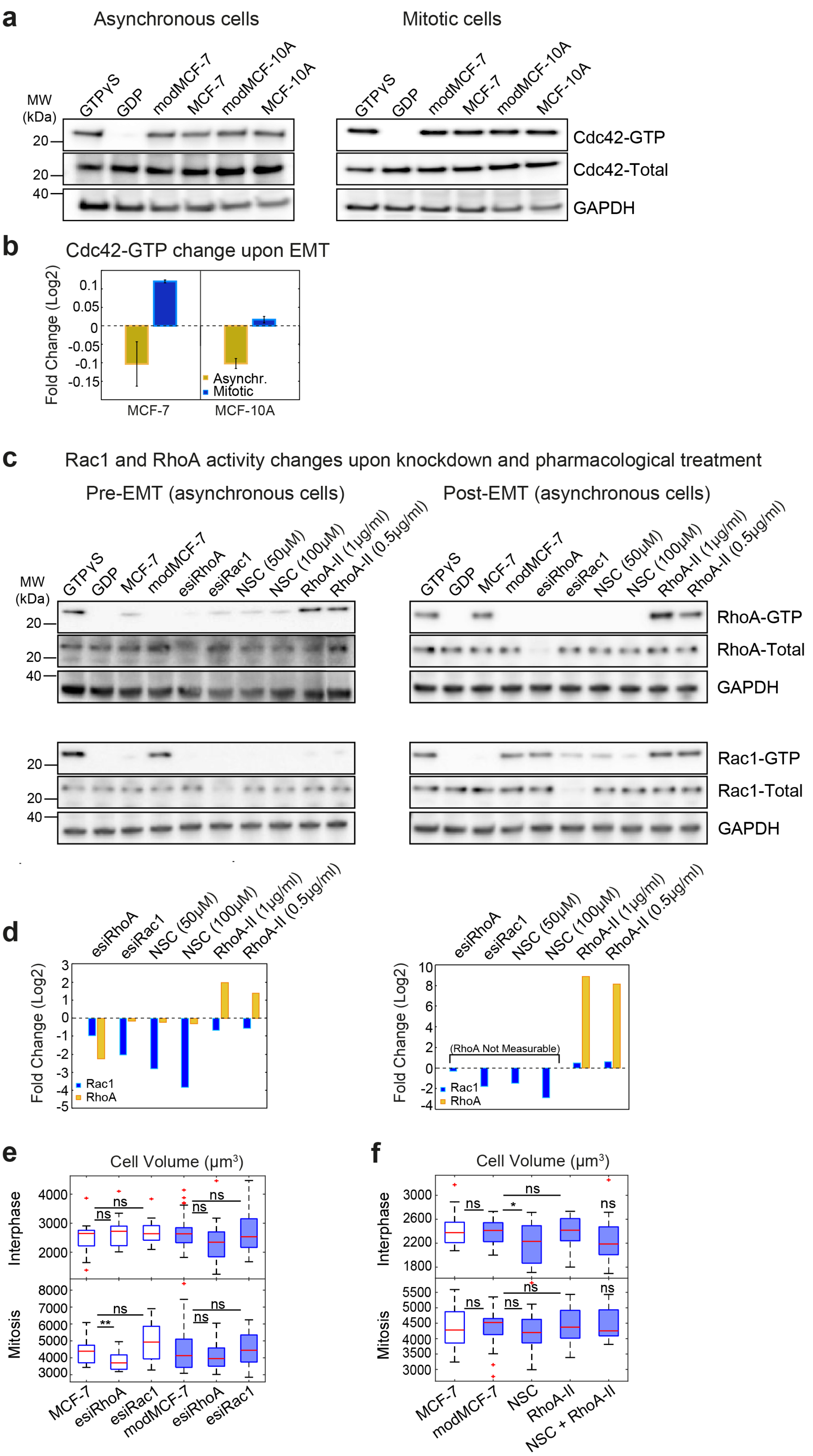

a
Asynchronous cells
Mitotic cells
MW (kDa)
GTPγS
GDP
modMCF-7
MCF-7
modMCF-10A
MCF-10A
Cdc42-GTP
Cdc42-Total
GAPDH
b
Cdc42-GTP change upon EMT
Fold Change (Log2)
Asynchr.
Mitotic
MCF-7
MCF-10A
c
Rac1 and RhoA activity changes upon knockdown and pharmacological treatment
Pre-EMT (asynchronous cells)
Post-EMT (asynchronous cells)
esiRhoA
esiRac1
NSC (50μM)
NSC (100μM)
RhoA-II (1μg/ml)
RhoA-II (0.5μg/ml)
RhoA-GTP
RhoA-Total
Rac1-GTP
Rac1-Total
d
(RhoA Not Measurable)
Rac1
RhoA
e
Cell Volume (μm³)
Interphase
Mitosis
ns
**
*
f
NSC
RhoA-II
NSC + RhoA-II

*Figure S4:* ***a-b,*** *Relative changes of active Cdc42 upon EMT. GTP-bound Cdc42 was pulled down from cell lysates using beads in pre and post-EMT conditions.* ***a,*** *Western blots showing active Cdc42 from cell lysates before and after EMT.* ***b,*** *Quantification of relative changes of Cdc42-GTP from Western blots (n=2).* ***c,*** *Western blots of pull-down assays (lysates from MCF-7 and modMCF-7 in asynchronous cell population) of Rac1-GTP and RhoA-GTP, and corresponding whole cell lysates of MCF-7 and modMCF-7 cells in control conditions and upon Rac1 or RhoA knock-down, 1 hours treatment with 50 μM or 100 μM NSC23766 and 2 hours treatment with 1 μg/ml or 0.5 μg/ml RhoA activator-II showing Rac1 and RhoA activity changes before and after EMT. Knockdown was achieved through RNA interference. esiFluc and esiKif11 were used as negative and positive controls in the experiment, respectively. Kif11 knock-down freezes cells in mitosis and generates a visible knock-down effect. Western blots clearly show that our knock-down protocol generates a successful knock-down of the target protein. The Western blots show successful activity changes of the target proteins upon knock-down or pharmacological treatments.* ***d,*** *Quantification of relative changes of Rac1-GTP and RhoA-GTP from Western blots in* ***c****. Quantifications were normalised by the GAPDH bands.* ***e,*** *Cell volumes in control conditions and after knock-down of Rac1 or RhoA corresponding to measurements presented in Fig. 3c-e, main text.* ***f,*** *Cell volumes in control conditions and after Rac1 inhibition or RhoA activation corresponding to measurements presented in Fig. 3f-h, main text (Post-EMT cells are referred to as modMCF-7 and mod-MCF-10A, respectively. . Number of cells measured: Panels* ***c-e****: Interphase: MCF-7 n=36, esiRhoA n=36, esiRac1 n=35, modMCF-7 n=31, esiRhoA n=26 and esiRac1 n=30, Mitosis: MCF-7 n=22, esiRhoA n=26, esiRac1 n=24, modMCF-7 n=28, esiRhoA n=28 and esiRac1 n=28. Panels* ***f-h****: Interphase: MCF-7 n=24, modMCF-7 n=24, NSC n=22, RhoA-II n=22 and NSC + RhoA-II n=24. Mitosis MCF-7 n=24, modMCF-7 n=28, NSC n=22, RhoA-II n=24 and NSC + RhoA-II n=22.* $^{ns}p > 0.05$, $^{*}p < 0.05$, $^{**}p < 0.01$, $^{***}p < 0.001$*).*

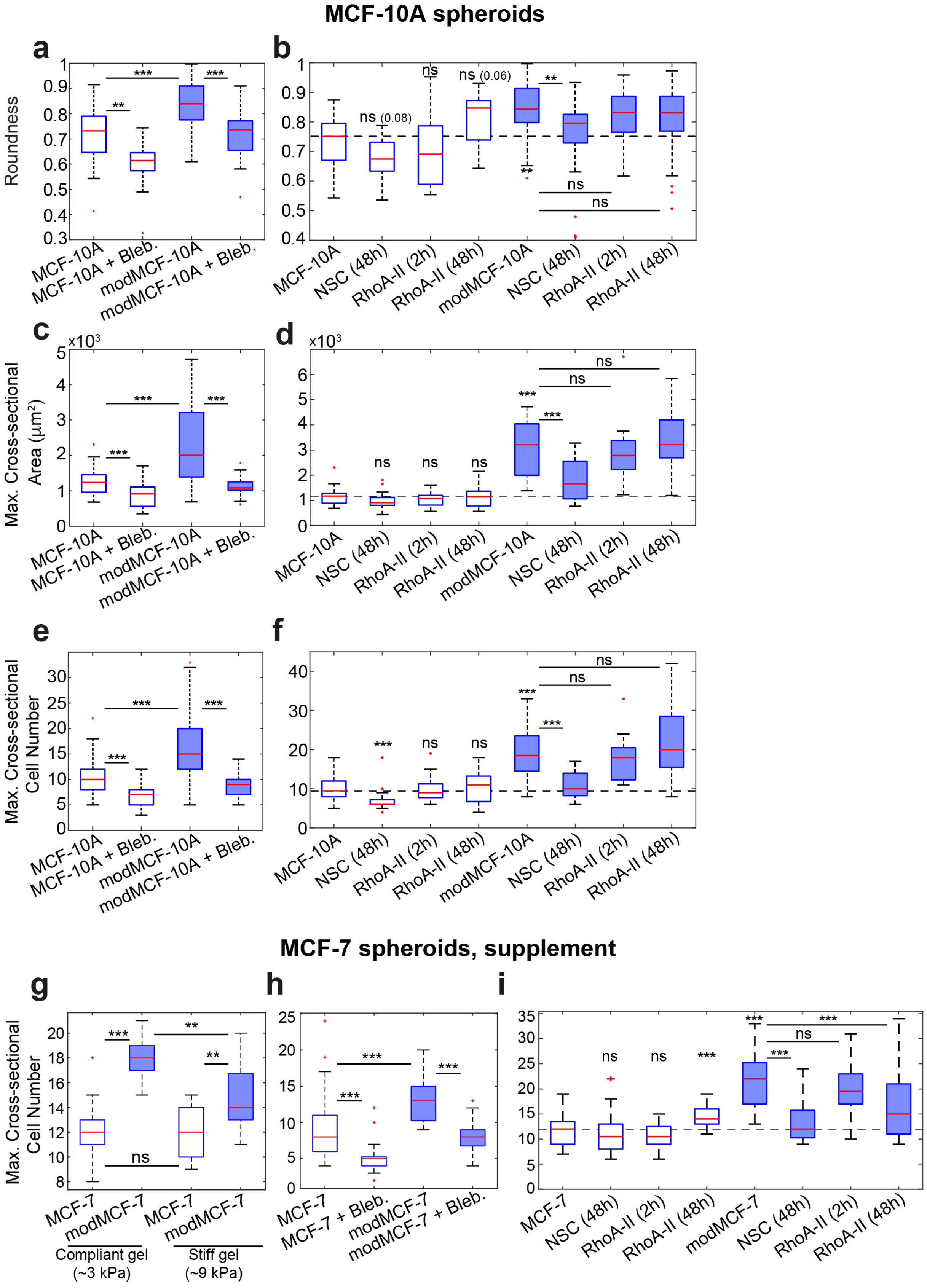


*Figure S5: The influence of EMT on mitotic roundness and spheroid size in MCF-10A tumor spheroids.* ***a,*** *Roundness of mitotic cells in compliant gels with or without inhibition of cortical contractility through the myosin inhibitor blebbistatin (10 µM).* ***b***, *Roundness of mitotic cells in compliant gels with*

*or without Rac1 inhibition upon 48 hours treatment with 50 µM NSC23766 (NSC) and RhoA activation upon 2 hours or 48 hours treatment with 1 µg/ml RhoA activator-II (RhoA-II).* ***c****, Influence of Myosin inhibition on spheroid size in compliant gels (Blebbistatin treatment for 2-days at 10 µM, see Materials and Methods) in pre- and post-EMT conditions.* ***d****, Influence of Rac1 inhibition on spheroid size upon 48 hours treatment with 50 µM NSC23766 (NSC) and RhoA activation upon 2 hours or 48 hours treatment with 1 µg/ml RhoA activator-II (RhoA-II) on spheroid size in pre- and post-EMT conditions.* ***e-f****, Number of cells in maximum cross-sectional area corresponding to* ***a-d.*** *).* ***g-i****, Number of cells in maximum cross-sectional area corresponding to Fig. 5f-h, main text. (Post-EMT cells are referred to as modMCF-7 and mod-MCF-10A, respectively.) Number of mitotic cells analyzed: Panel* ***a****: MCF-10A n=27, MCF-10A+Bleb., n=10, modMCF-10A n=58 and modMCF-10A+Bleb. n=27. Panel* ***b****: MCF-10A n=14, NSC (48h) n=14, RhoA-II (2h) n=10, RhoA-II (48h) n=13, modMCF-10A n=37, NSC (48h) n=33, RhoA-II (2h) n=38 and RhoA-II (48h) n=96. Panel* ***c****: MCF-10A= 41, MCF-10A+Bleb. n=39, modMCF-10A n=40, modMCF-10A+Bleb. n=35. Number of spheroids analyzed: Panel* ***d****: MCF-10A n=20, NSC (48h) n=23, RhoA-II (2h) n=21, RhoA-II (48h) n=21, modMCF-10A n=20, NSC (48h) n=19, RhoA-II (2h) n=15 and RhoA-II (48h) n=19. Panel* ***e****: MCF-10A= 41, MCF-10A+Bleb. n=39, modMCF-10A n=40, modMCF-10A+Bleb. n=35. Panel* ***f****: MCF-10A n=20, NSC (48h) n=23, RhoA-II (2h) n=21, RhoA-II (48h) n=21, modMCF-10A n=20, NSC (48h) n=19, RhoA-II (2h) n=15 and RhoA-II (48h) n=42. Panel* ***g****: MCF-7 n=37, modMCF-7 n=34, (Stiff gel) MCF-7 n=17 and modMCF-7 n=15. Panel* ***h****: MCF-7= 26, MCF-7+Bleb. n=25, modMCF-7 n=31, modMCF-7+Bleb. n=33. Panel* ***i****: MCF-7 n=40, NSC (48h) n=40, RhoA-II (2h) n=40, RhoA-II (48h) n=31, modMCF-7 n=41, NSC (48h) n=39, RhoA-II (2h) n=40 and RhoA-II (48h) n=42.* $^{ns}p > 0.05$, $^{*}p < 0.05$, $^{**}p < 0.01$, $^{***}p < 0.001$*).*

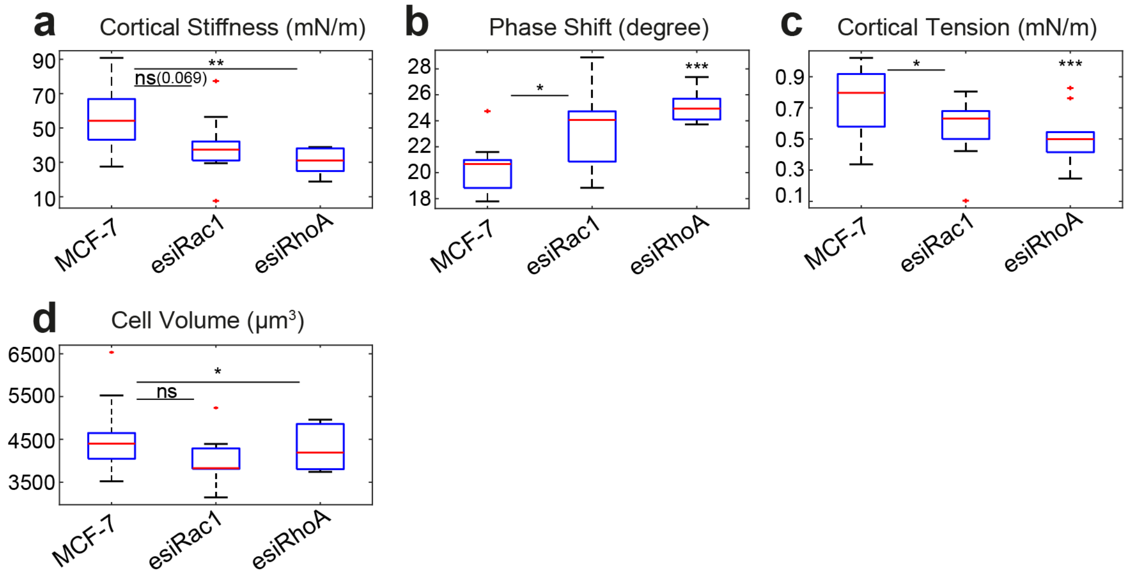


*Figure S6: Cortex mechanics of metaphase MCF-7 cells (without STC) before and after Rac1 and RhoA knock-down. Cells were identified by roundness and presence of a metaphase plate (Number of cells analyzed: MCF-7 n=15, esiRac1 n=11, esiRhoA n=10. Measurements are representative for two independent experiments.* $^{ns}p > 0.05$, $^{*}p < 0.05$, $^{**}p < 0.01$, $^{***}p < 0.001$*).*

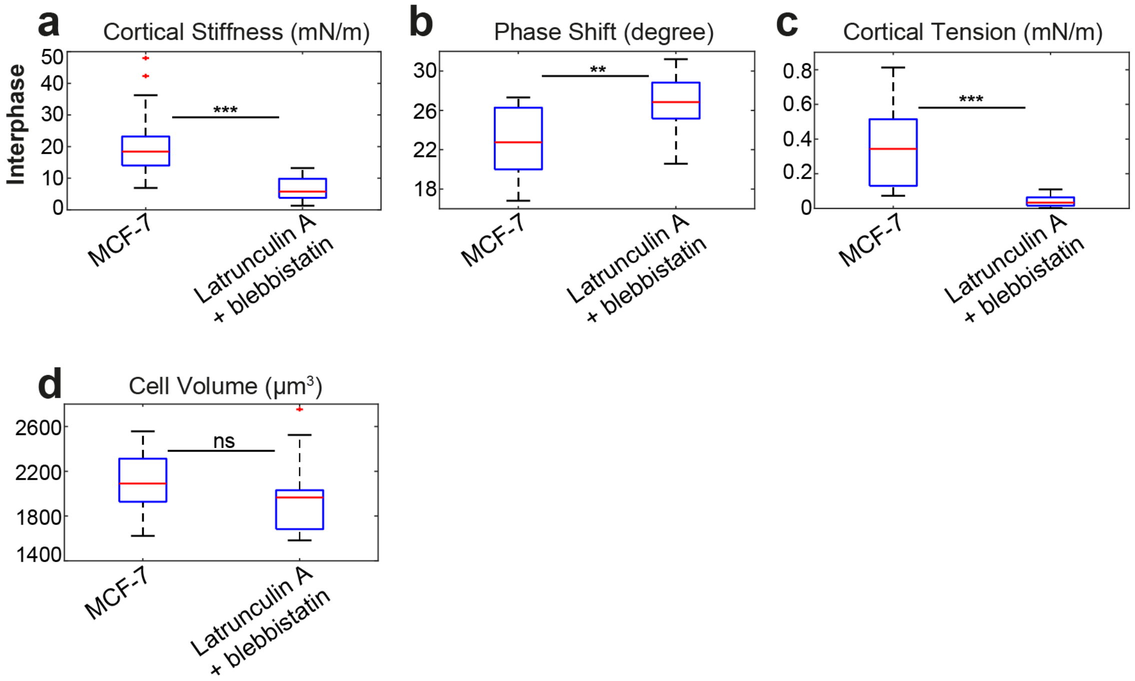


*Figure S7: Cortex mechanics of interphase MCF-7 cells before and after 30 minutes incubation with 200 nM latrunculin A & 10 μM blebbistatin. (Number of cells analyzed: MCF-7 n=20, Latrunculin A + blebbistatin n=20. Measurements are representative for two independent experiments. $^{ns}p > 0.05$, $*p < 0.05$, $**p < 0.01$, $***p < 0.001$).*
